# Ongoing surveillance protects tanoak whilst conserving biodiversity: applying optimal control theory to a spatial simulation model of sudden oak death

**DOI:** 10.1101/773424

**Authors:** E.H. Bussell, N.J. Cunniffe

## Abstract

The sudden oak death epidemic in California is spreading uncontrollably. Large-scale eradication has been impossible for some time. However, small-scale disease management could still slow disease spread. Although empirical evidence suggests localised control could potentially be successful, mathematical models have said little about such management. By approximating a detailed, spatially-explicit simulation model of sudden oak death with a simpler, mathematically-tractable model, we demonstrate how optimal control theory can be used to unambiguously characterise effective time-dependent disease management strategies. We focus on protection of tanoak, a tree species which is culturally and ecologically important, but also highly susceptible to sudden oak death. We identify management strategies to protect tanoak in a newly-invaded forest stand, whilst also conserving biodiversity. We find that thinning of bay laurel is essential early in the epidemic. We apply model predictive control, a feedback strategy in which both the approximating model and the control are repeatedly updated as the epidemic progresses. Adapting optimal control strategies in this way is vital for effective disease management. This feedback strategy is robust to parameter uncertainty, limiting loss of tanoak in the worst-case scenarios. However, the methodology requires ongoing surveillance to re-optimise the approximating model. This introduces an optimal level of surveillance to balance the high costs of intensive surveys against improved management resulting from better estimates of disease progress. Our study shows how detailed simulation models can be coupled with optimal control theory and model predictive control to find effective control strategies for sudden oak death. We demonstrate that control strategies for sudden oak death must depend on local management goals, and that success relies on adaptive strategies that are updated via ongoing disease surveillance. The broad framework allowing the use of optimal control theory on complex simulation models is applicable to a wide range of systems.

## Introduction

Sudden oak death (SOD), caused by the oomycete *Phytophthora ramorum*, affects a broad range of hosts including oaks, tanoak and bay laurel. The disease was first detected in California in 1995 and has significantly impacted the nursery trade, and devastated populations of coast live oak and tanoak in California. SOD is currently found in areas covering over 2,000 km^2^ in California [1], with an estimated $135M loss in property values attributed to the disease [2]. Landscape scale mathematical models have shown that widespread eradication would require an infeasibly large removal of trees, making it now impossible [3]. Control has been shown to be effective at smaller scales though. Management of the isolated outbreak in Oregon has been effective at slowing the spread, containing the disease within Curry county [4]. In some locations, control in Oregon has shown that local eradication, whilst difficult, is possible [5]. Mathematical models however, have had little to say about how this local control should be optimised.

Time-varying controls can be optimised using branch of applied mathematics called optimal control theory (OCT) [6]. However, its application is often limited to relatively simple models of disease spread for which the underpinning analysis remains tractable [7]. Previous work has shown a proof of concept application of OCT to a more complex model using an approximate model for optimisation [8]. OCT can be applied to the approximate model, and the resulting optimal control applied to the simulation model using one of two frameworks: open-loop or model predictive control (MPC) [9]. Whereas the open-loop framework applies the optimal control to the simulation model for all time, MPC allows for feedback from the simulation through ongoing surveillance and re-optimisation of the control at regular update times. Both frameworks allow optimisation of a complex system, and show how mathematical models can be used to optimise practical disease management: methods that could in principle be applied to optimise local control of SOD (or any other disease).

One host particularly impacted by SOD is tanoak, *Notholithocarpus densiflorus*, a medium sized Californian tree related to the American chestnut. Tanoak trees are highly valued by Native American tribes in Northern California for their acorns, which are used for trade and are an important source of food [10]. Tanoak is also believed to be highly important to coast redwood forest ecosystems [11, 12]. The spread of *P. ramorum* is a significant threat to tanoak. If decline in populations continues, extinction of this important species is possible [13]. Tanoak trees of all ages are highly susceptible to SOD and have a very high mortality from the disease [14, 15], with larger trees that produce more acorns disproportionately affected [16]. Because of the importance of retaining tanoak, management strategies should be designed to protect this vital species.

The presence of California bay laurel (*Umbellularia californica*) is a key factor in the decline of tanoak due to SOD. *P. ramorum* can sporulate prolifically on bay trees, but that host is not killed by the disease [17]. Field studies have shown that removal of bay laurel, and thinning of susceptible trees to reduce host density, can control disease spread [18]. Environmental niche models have been developed to classify SOD invasion risk [19, 20], and dynamic mathematical models have been used to predict SOD spread at large scales [21, 22]. Only one model captures dynamics at a small enough scale to show the decline of overstorey tanoak at the forest stand level (∼20 ha) [16]. Studies have used this model to demonstrate the effectiveness of one-off interventions [18, 23], but do not assess time-dependent controls, and strategies have not been optimised in order to protect tanoak.

Whilst a clear objective for control is to protect tanoak, this must not be at the cost of the overall health of the forest. Trees in forests are important for wildlife habitats and food sources, recreational uses and carbon fixation [24], and maintaining diversity ensures that these and other important ecosystem services are provided [25, 26]. Beyond this, diverse forests are more resilient to other disease threats [27]; there is little point to a control strategy that protects tanoak from SOD but makes the forest vulnerable to attack from another pathogen. The management goal must therefore capture a balance between protection of tanoak, and continued host diversity for provision of ecosystem services. The balance between these two objectives however, will depend on the overall local management goals for the forest stand in question, and economic valuation of the ecosystem services [28].

In this paper we will optimise SOD control strategies to protect tanoak within individual 20 ha mixed species forest stands, whilst also conserving biodiversity. We will show how time-dependent control strategies can be found using OCT on an approximate model of disease dynamics, and applied to a complex spatial simulation model. By comparing the open-loop and MPC frameworks, we demonstrate the importance of ongoing disease surveillance for effective control, and the importance of re-optimising and adapting the management strategy. Most importantly, we will look at the effects of parameter and observational uncertainty on control efficacy, testing under what conditions and to what extent ongoing surveillance is necessary.

We seek to answer the following key questions:

1. How can the open-loop and MPC frameworks be used to allow the insights from OCT to be applied to a detailed spatially-explicit simulation model of SOD?
2. How should time-dependent controls be deployed under resource constraints to best preserve the valuable tanoak population in coastal redwood forests, and how do these strategies compare with current recommended practice?
3. How robust and reliable are these control results? In particular, how important is the ongoing surveillance of disease progression and re-optimisation of control when there is parameter uncertainty and imperfect sampling?

## Results

We use a spatially-explicit simulation model of SOD spread and tanoak population decline, adapted from a model by Cobb *et al*. [16] to include more realistic inoculum dispersal (S1 Text). The model tracks the stem density and disease progression in 400 cells across a 20 ha forest stand (Fig 1D–1E) of redwood, which is epidemiologically inactive, bay laurel which is highly infectious but does not die from the disease, and tanoak. Tanoak is tracked in four age classes, allowing analysis of overstorey tanoak decline (Fig 1A). Optimisation of time-varying control using the full spatial simulation is impossible, so a non-spatial approximate model of the dynamics is used and the optimal control strategy applied to the simulation using the open-loop and MPC frameworks. The approximate model was fitted to the simulation model and closely matches the progression of disease (Fig 1B–1C).

**Figure 1:**
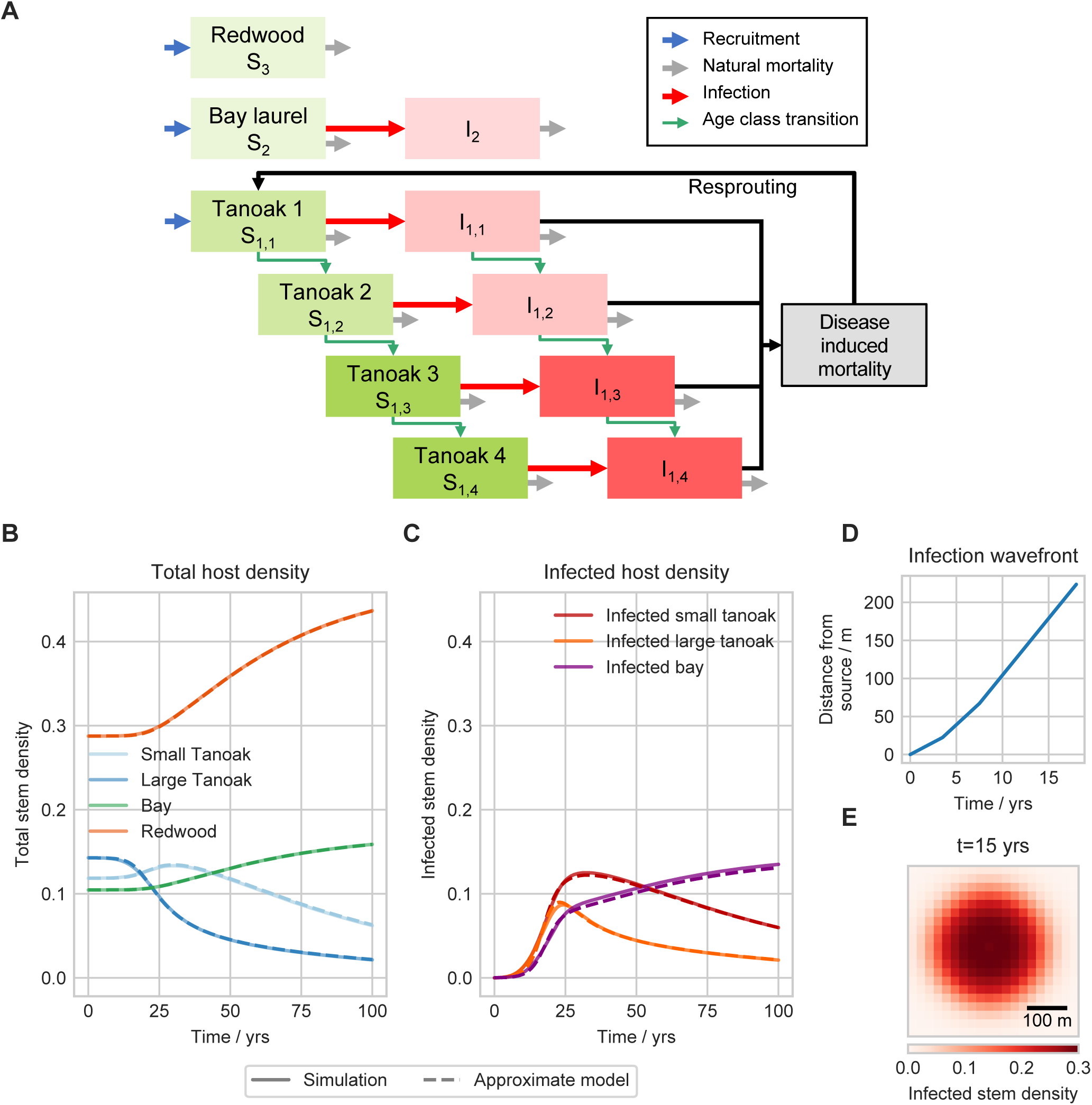
Stand level simulation and approximate models. A: The possible host states and transitions in both the simulation and approximate models. Only bay and tanoak are epidemiologically active, with the tanoak age classes grouped into small (tanoak 1 and 2) and large (tanoak 3 and 4) categories. We here focus on protection of large (overstorey) tanoak. B: The overall stem density for each species class under no control, with the dashed lines showing the approximate model. The dynamics of the fitted approximate model closely match the spatial simulation dynamics, showing significant decline of overstorey tanoak. C: The disease progress curves track the infected stem density. The approximate model is fitted by matching the disease progress curves between the two models. As the host demography is the same in the simulation and approximate models, this leads to the overall host dynamics closely matching, as shown in B. D: Following introduction at *t* = 0 to the centre cell, the distance from the source to the epidemic wavefront, here defined as the furthest cell with an infected stem density above 0.05. After a short transient (∼5 years to 10 years), the epidemic wave travels at a constant speed. E: The spatial distribution of infected hosts in the simulation model, 15 years after introduction.

### Surveillance and MPC improve disease management

Time-dependent roguing (i.e. removal of infected hosts), thinning (i.e. removal of hosts of all infection statuses) and protectant application (which reduces tanoak susceptibility by 25 %) controls are optimised using the approximate model. Controls are deployed in stages, with controls kept constant over each 5 year stage. The controls are optimised with an objective to maximise retention of healthy large tanoak after 100 years, alongside conserving biodiversity over that time. The optimal strategy in the approximate model thins bay laurel early in the epidemic, then switches to thinning redwood before using the protectant methods once sufficient thinning has been completed. Roguing of both tanoak and bay is carried out to varying extents throughout the epidemic (Fig 2A). Using the open-loop framework, the optimal strategy is lifted to the simulation, and applied to the spatial model for the entire time. In the simulation model the open-loop strategy slows tanoak decline, retaining tanoak in the forest stand after 100 years (Fig 2B). However, the non-spatial approximation cannot match the simulation model over the entire epidemic and there is significant tanoak decline towards the end of the time period caused by a late re-emergence of disease.

**Figure 2:**
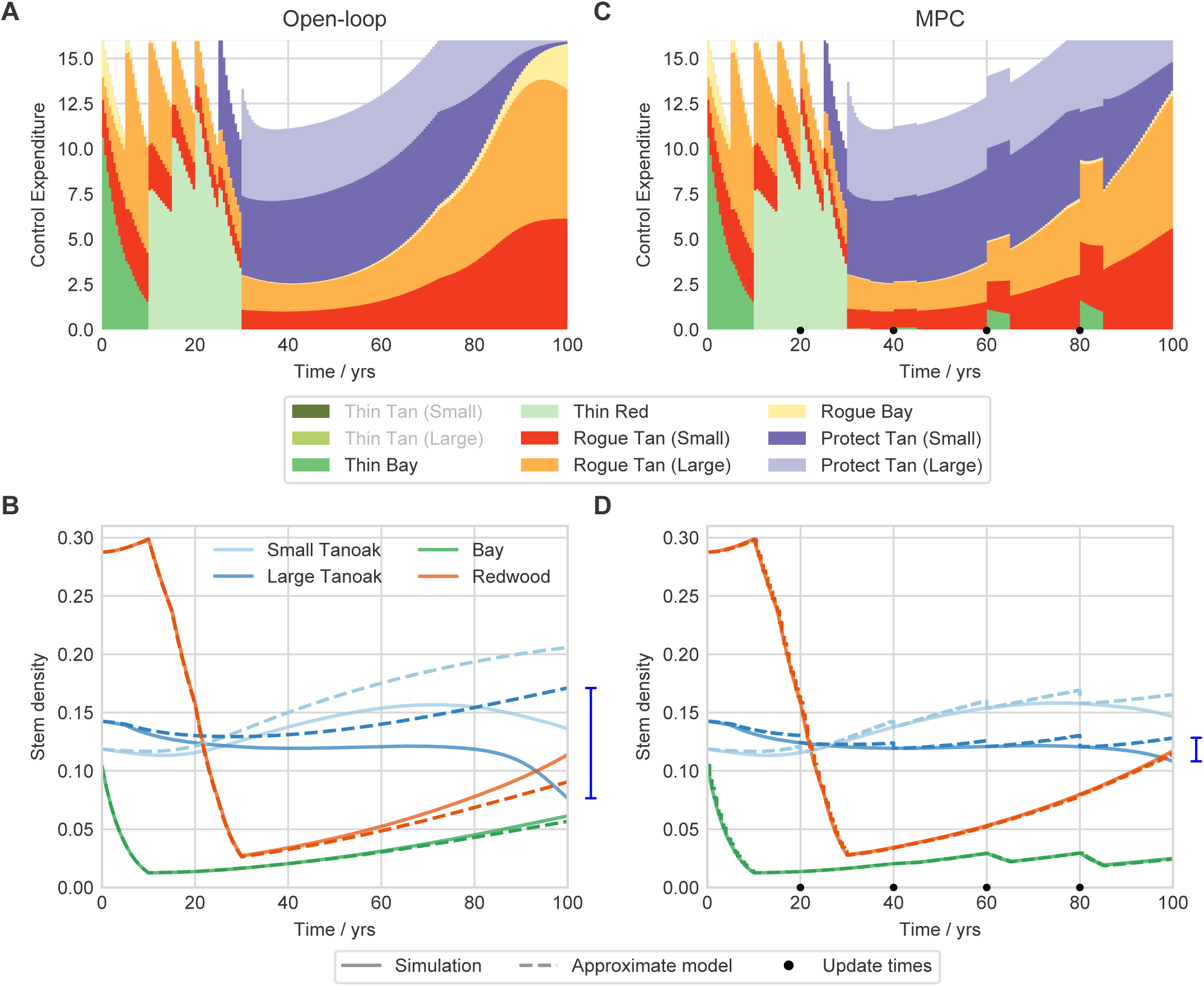
MPC improves protection of tanoak over the open-loop framework. The optimised control strategy in the spatial simulation model is found using optimal control theory on the simpler, non-spatial approximate model. This strategy is then applied directly to the spatial simulation model. A and B show open-loop control, C and D show MPC where the approximate model is reset and re-optimised at regular update times. Tanoak age classes 1 and 2 are grouped as small, and 3 and 4 as large tanoak. A: The resources allocated to each control method over time using the open-loop framework. Control proportions (*f*_*i*_ in S1 Text) are fixed over 5 year intervals, but as the number of hosts changes during each interval, this leads to variable expenditure over the 5 years. Greyed out control methods in the legend are not used in either strategy, as OCT identifies them as ineffective. Heavy thinning of bay is carried out early in the epidemic followed by thinning of redwood. Resources are allocated to roguing throughout, but protectant controls are only used once resource intensive thinning has been completed. B: The host dynamics corresponding to the open loop control in A, for both the approximate and simulation models. The approximation degrades towards the end of the epidemic, leading to unanticipated tanoak decline in the simulation. The blue bar highlights the difference between the approximate and simulation models in the density of large tanoak. C: The MPC resource allocation, updated every 20 years, shows a similar pattern to the open-loop strategy. However, additional thinning is carried out after each update later in the epidemic to manage the bay population, within which infection has increased more than anticipated by the approximate model. D: The corresponding host dynamics show that MPC repeatedly resets the approximate model trajectory, allowing more informed control decisions. This minimises the tanoak decline seen using open-loop, and gives a much lower error in the estimate of large tanoak stem density (blue bar).

An alternative strategy is to use the MPC framework, in which the approximate and simulation models are run concurrently. At regular update times the approximate model is reset to match the current state of the simulation model, and the control is re-optimised. This ensures the approximate model closely matches the simulation, allowing for more informed optimal control strategies. These updated controls are then lifted to the simulation model going forward until the next update time, here every 20 years. Applying this strategy results in additional thinning of bay after each update time (Fig 2C). This is to stem increased infection in bay, unanticipated in the approximate model because of overoptimistic assessment of the control performance. The MPC strategy keeps the late disease re-emergence under control. Through the continued surveillance of the simulation, the approximate model is kept more closely aligned with the simulation model. This makes the control more effective at slowing tanoak decline (Fig 2D).

The MPC strategy retains more tanoak than the open-loop strategy, primarily due to increased thinning of bay laurel. This additional thinning results in lower biodiversity conservation than the open-loop framework (Fig 3A), but this is balanced by significantly more healthy large tanoak after 100 years (Fig 3B). When compared with the situation under no control interventions, both strategies are highly effective at delaying tanoak decline. The overall objective is a combination of tanoak retention and biodiversity conservation, and whilst both methods outperform no control, the continued surveillance in the MPC framework gives the most effective disease management. The benefit of slowing tanoak decline is not captured fully by the objective function though, and open-loop is successful at delaying impacts on tanoak.

**Figure 3:**
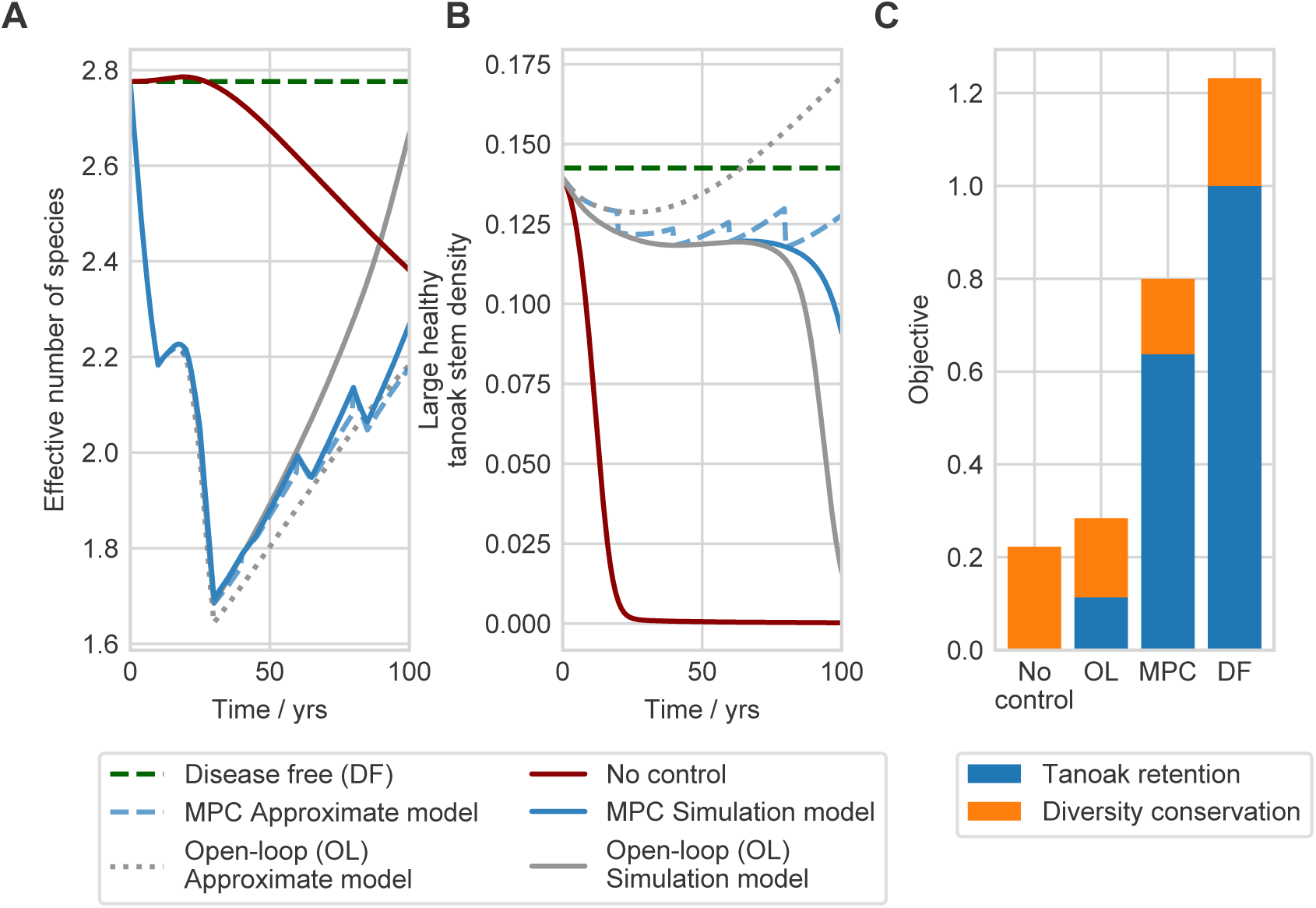
Comparing the open-loop and MPC frameworks. A: The effect of control on the diversity in the forest stand, shown as the effective number of equally-common species. Any control — be it optimised via open-loop or MPC — is damaging to diversity, but MPC in fact has a slightly larger impact. For MPC the simulation and approximate model dynamics are shown, with the approximate model resetting every 20 years. B: The stem density of healthy large tanoak over time, showing significant decline under no control. The open-loop strategy slows tanoak decline, but MPC is more effective. C: The overall performance of the strategies as measured by the objective function. MPC is slightly more damaging to diversity than open-loop, but this is balanced by retaining significantly more healthy tanoak.

### How does conserving biodiversity affect tanoak retention?

The objective we have used to optimise control incorporates both tanoak retention and biodiversity conservation. How does including biodiversity in the management goal affect the optimal control strategy and the resulting disease management? When biodiversity is not included in the objective, the optimal strategies under both the open-loop and MPC frameworks result in complete removal of the bay and redwood populations (Fig 4A and S1 Figure). For many forest managers and conservationists this would be an unacceptable cost to slow decline of a single important species [11]. Including the many benefits of a diverse forest stand in the objective function ensures that control retains all species, but does result in less effective retention of tanoak, at least when using the open-loop framework (Fig 4B). Whilst including biodiversity in the open-loop framework results in the functional extinction of tanoak (i.e. no ecosystem services are provided), the MPC framework can still retain healthy overstorey tanoak. The continued surveillance and re-optimisation allows the control to manage the disease whilst also conserving biodiversity, even for much higher biodiversity benefits than are used by default (Fig 4C–4D).

**Figure 4:**
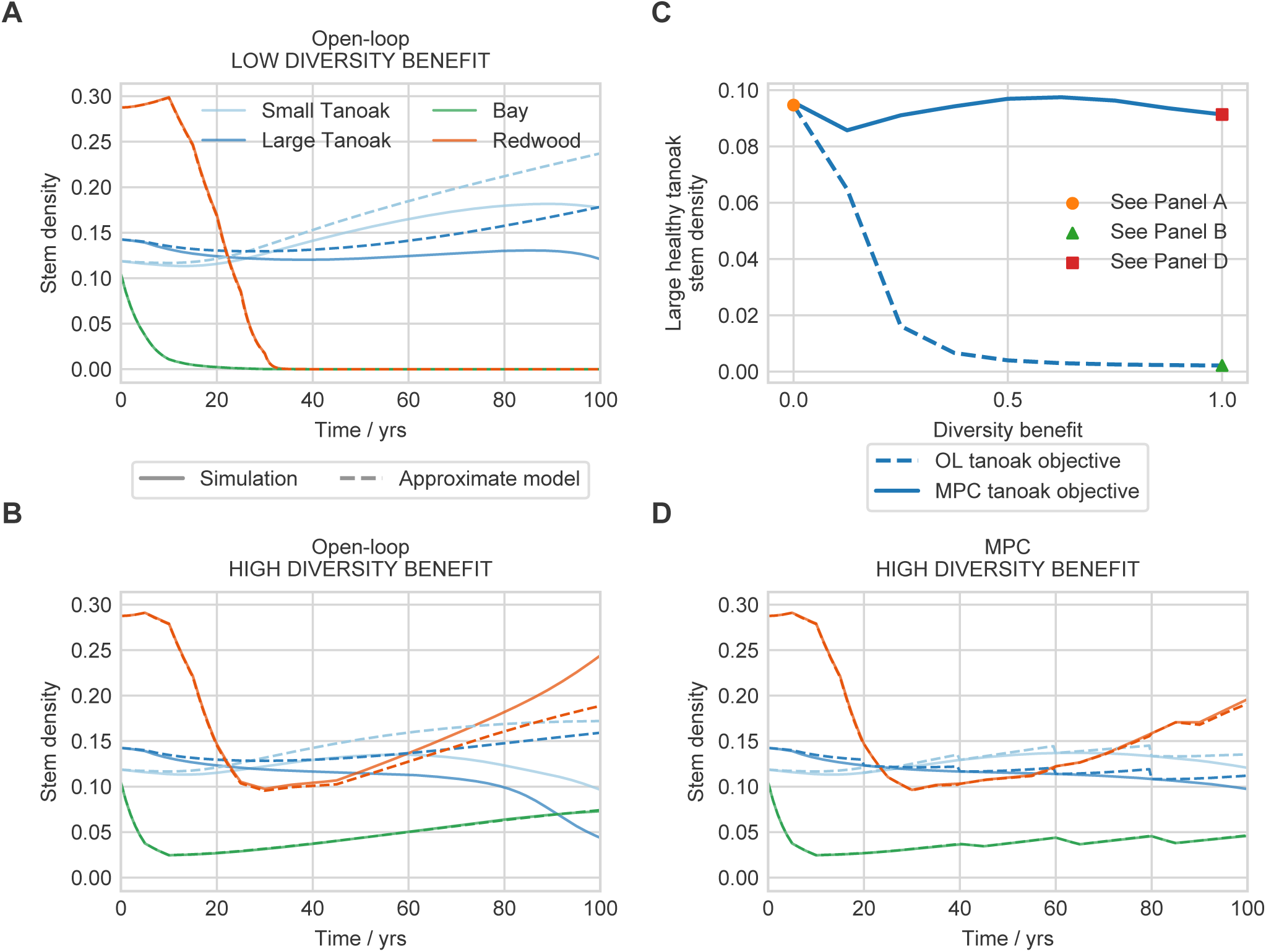
Impact of the diversity objective on the optimal control strategy. Including a diversity metric in the objective function is necessary to ensure protection of tanoak is not carried out to the complete detriment of the rest of the forest. A: With no benefit to diversity the optimal control (using the open-loop framework) removes all bay and redwood hosts. A similar strategy is seen using MPC (S1 Figure). B: High values of the diversity benefit lead to control that does not stop tanoak decline, at least in the open-loop case. C: The healthy tanoak retained in the forest decreases as diversity becomes more important in the open-loop case. Note that the y axis here is healthy large tanoak, so does not show the same stem density as in A and B where infected hosts are also counted. For the MPC framework however, the control can conserve biodiversity whilst also protecting tanoak. The precise performance depends on the fit of the approximate model under each control strategy, leading to above or below expected performance for some values of the biodiversity benefit. For reference, the baseline value of the diversity benefit used throughout the paper is 0.25. D: The host density with high diversity benefit under MPC shows that bay and redwood can be retained whilst also limiting tanoak decline.

### MPC is robust under uncertainty

#### Uncertainty in infection rate parameters

In reality, precise values of infection rates are never known. These parameters are often fitted to limited data with Bayesian techniques, giving a probability distribution of values [e.g. 29–32]. We tested how the open-loop and MPC frameworks handle this type of uncertainty in the system dynamics. Uncertainty is introduced by sampling values from a distribution of infection rates for each species in the simulation model. The approximate model is re-fitted to this ensemble of 200 simulation runs (Fig 5A–5B). To test control on these parameter distributions, for a single draw of infection rates from the distribution, the fitted approximate model is used to run the open-loop and MPC frameworks. This is repeated for 200 draws from each distribution.

**Figure 5:**
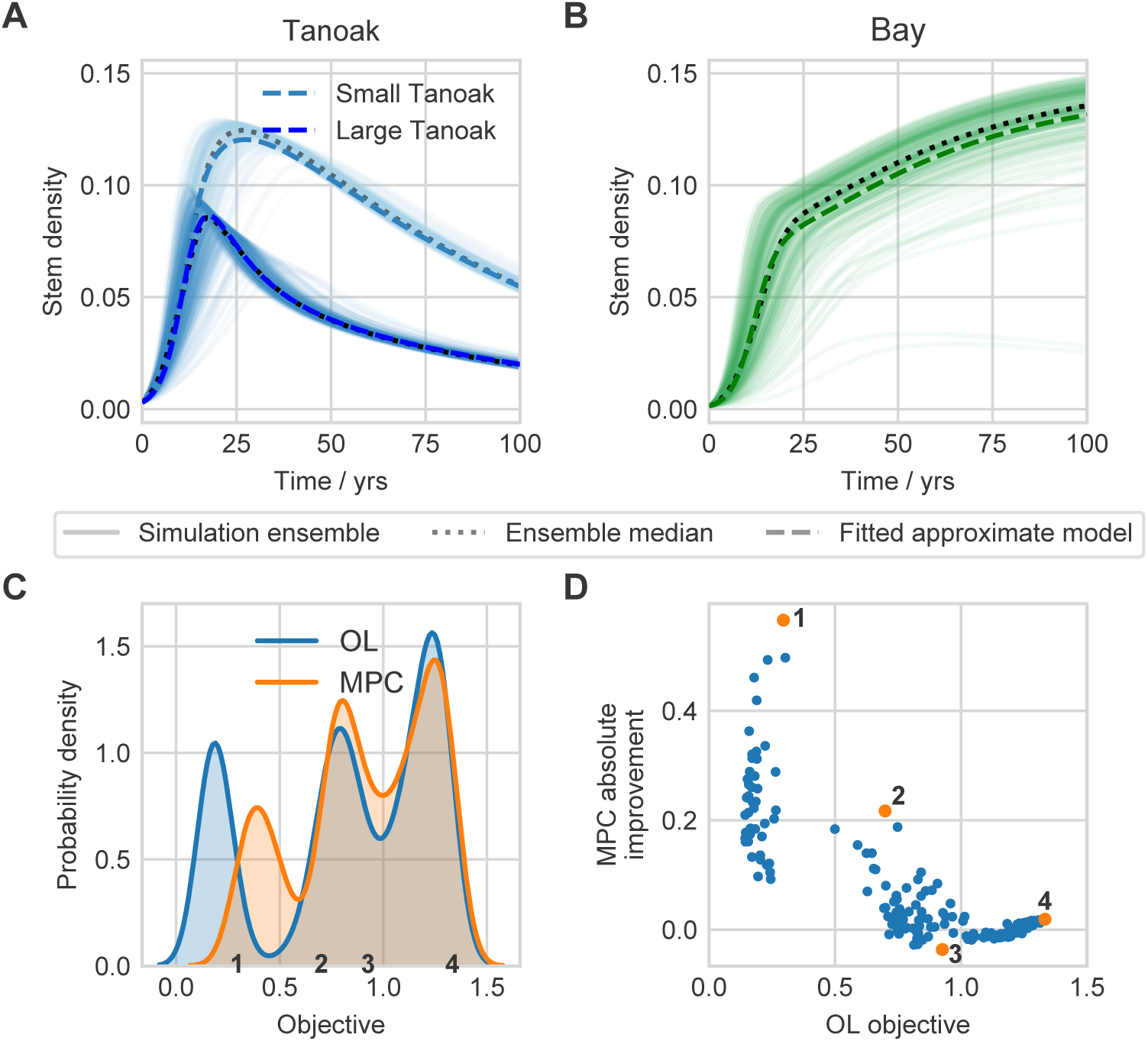
Effect of parameter uncertainty on control performance. The approximate model is fitted to an ensemble of simulation runs without control, with infection rate parameters drawn from a truncated normal distribution. A: The ensemble and fitted tanoak dynamics. B: The ensemble and fitted bay dynamics. C: The distribution of objective values using open-loop and MPC across 200 draws of simulation parameters. D: The absolute improvement of the MPC strategy over open-loop, as a function of the open-loop objective. The MPC framework performs well in the worst-case scenarios, improving control to the largest extent when open-loop performs badly. Four individual cases have been highlighted in panels C and D. Further details for each of these, highlighting what drives the differences in performance, are shown in S4 Text.

The distribution of the resulting objective values under the open-loop and MPC frameworks shows that MPC improves the worst-case scenarios, i.e. the MPC updates are most beneficial when the disease is hardest to manage (Fig 5C). The open-loop framework gives a distribution of objectives with a worse minimum value than MPC. The continued surveillance of MPC generally improves control, but the greatest improvements are seen when the epidemic is difficult to control, making the open-loop framework ineffective (Fig 5D). When objective values are high, and so the epidemic is easy to control, there is little difference between open-loop and MPC.

#### Imperfect sampling

The re-optimisation of control in the MPC framework requires accurate information about the current state of the forest at each update step, i.e. every 20 years by default. However, full forest stand surveys are expensive. We tested whether the intensity of these update surveys can be reduced whilst maintaining effective control, and whether open-loop strategies can ever outperform MPC with low quality surveillance. Imperfect sampling is introduced at each MPC update step, by sampling only a proportion of the forest stand giving imperfect observation of the state of the epidemic. This imperfect observation is then fed back into the approximate model, with control re-optimised in MPC based on this imperfect knowledge.

As the proportion of area sampled decreases, the uncertainty in the outcome of MPC increases (Fig 6A). The median performance of MPC also decreases. This is because as less of the forest is sampled, there is a higher chance that infected hosts will be missed during surveillance, and so the rate of disease spread will be underestimated. As the proportion of the stand that is surveyed increases, so too do the surveillance costs (Fig 6B). The disease costs however, reduce with more tanoak retained as a result of more informed and effective control strategies. Balancing these two costs results in an optimal level of surveillance effort (Fig 6C). The precise location of this optimum depends on the balance between tanoak retention, biodiversity conservation, and surveillance costs: a decision that must be made in the context of local forest management goals. It is clear though, that some level of continued surveillance and re-optimisation through the MPC framework is necessary for effective control.

**Figure 6:**
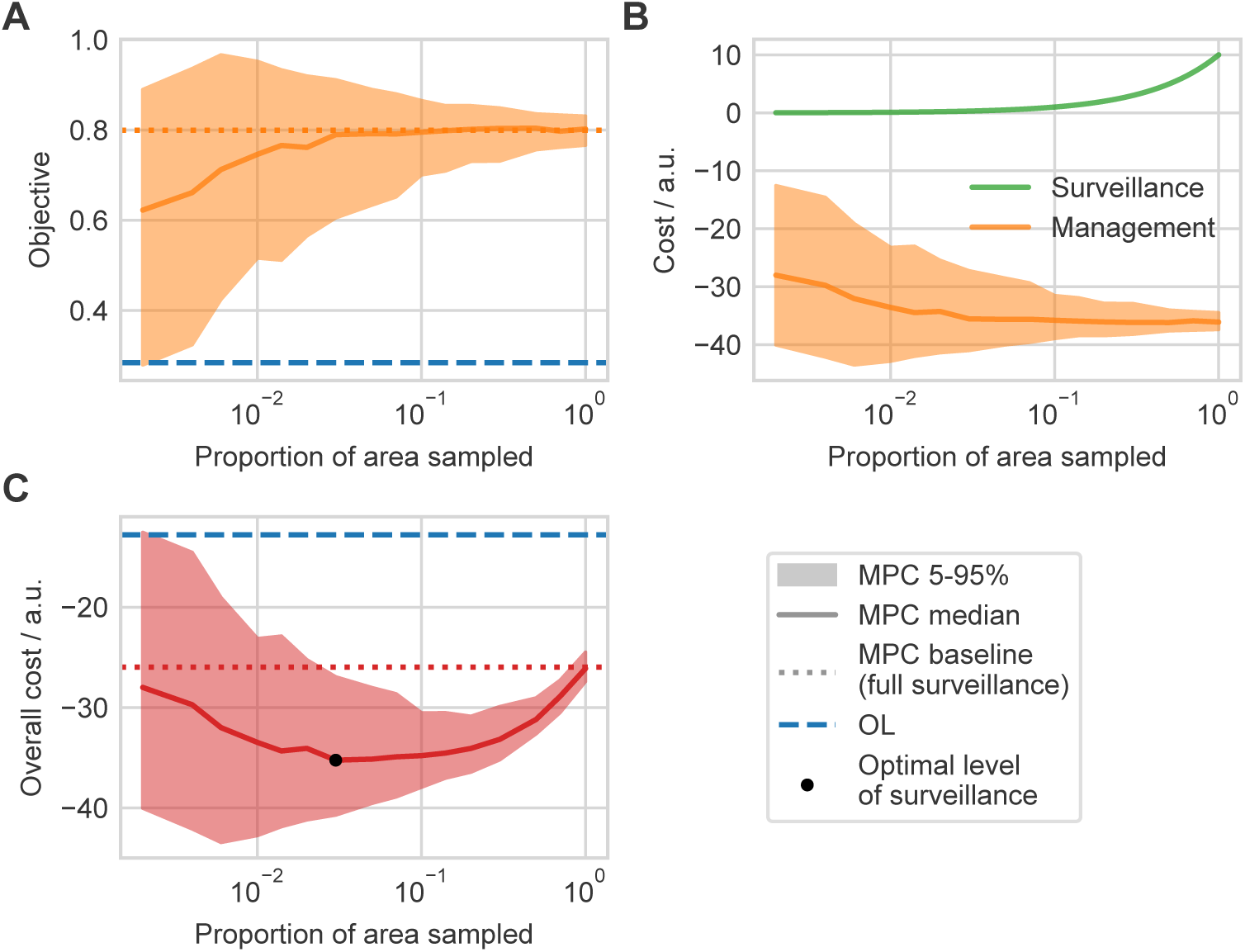
Effect of imperfect surveillance on control performance. Random sampling of a proportion of the forest stand at each MPC update step leads to imperfect surveillance. A: The distribution of objective values as a function of the proportion of the stand surveyed at MPC updates. Imperfect surveillance leads to reduced performance on average. In some cases over-estimation of the level of infection leads to improved control by chance. In general, over-estimation is ineffective because of unnecessary biodiversity losses. B: The costs of surveillance and benefits of improved management. More intensive surveillance is more expensive, but improves average performance. The management benefits are given by the objective multiplied by a negative scaling factor, here −45. The surveillance costs are fixed at 10 units per survey, multiplied by the proportion of the forest that is being sampled C: The total costs of surveillance and benefits of management. Balancing the costs and benefits leads to an optimal level of surveillance that depends on how management benefits are valued compared to the surveillance costs.

## Discussion

Despite the empirical effectiveness of local SOD management strategies [5], mathematical modelling studies have had little to say about how local control should be optimised. Poorly designed disease management strategies can lead to expensive failures of control, as was the case for control of Dutch elm disease in the UK [33] and citrus canker in Florida [34]. Although detailed simulation models have been used to optimise management strategies for a number of plant pathogens [3, 32, 35–41], computational constraints mean that strategies under test are typically restricted to one from a small set of possibilities in which the amount of control remains constant over time. Here we have shown how frameworks for using optimal control theory [9] can be applied to realistic simulation models to understand a practical disease management question, and identify effective management strategies. By comparing the open-loop and MPC frameworks, we demonstrated that continued surveillance and re-optimisation of the strategy improves management, and leads to control that is robust to parameter uncertainty and imperfect sampling. Current advice for management of SOD centres around removal of the spreader species, bay laurel, as well as infected tanoak and bay, and susceptible tanoak close to known infections [42]. Application of protective chemical treatments is also recommended for high value trees that are close to infections and known not to be currently infected [43]. The strategies found here using OCT are broadly similar in nature to these recommendations, but include significant time dependence, i.e. dependence on the state of the local epidemic. We have found that thinning of bay laurel is very important to the success of disease management, echoing results of previous modelling studies [18, 44]. Management advice from the US Forest Service [24] also suggests removal of bay trees, and even complete area-wide removal of bay in some cases. We found that roguing of tanoak is more important than roguing of infected bay, as thinning would reduce the bay population density sufficiently for disease control. Our results show that with continued surveillance and careful optimisation of controls, disease can be successfully managed. This can be done whilst maintaining a bay population which may be ecologically important. However, when biodiversity benefits are less important, control is always easier and more effective with additional removal of bay laurel.

Application of chemical protectants is only recommended in practice for individual high value trees close to known infections [42]. The strategies we have found deploy significant protection resources though. In fact, the protectant application only has a very minor effect on the performance of the strategies (S3 Text), and is unlikely to be cost-effective. Our formulation of the budget constraint implements a maximum expenditure where a fixed amount of money is put aside for SOD control, rather than minimising total costs. This captures governmental allocation of money for SOD control. In our model, when the optimal levels of roguing and thinning do not use the entire budget, the surplus can be allocated to protectant application. In practice though, control methods will also be individually assessed for cost-effectiveness, and given the limited effect of the protectant strategies it seems unlikely that they would be used.

A possible limitation of the strategies we have found is their complexity. Control inputs were held constant over 5 year stages so that resources do not have to be continually moved, but the strategies are nevertheless still complex in their time dependence and relative allocations to multiple control methods. However, the findings of our results could still be useful. Building up an intuition about what drives the optimal strategy could lead to more practical advice. For example, in the open-loop and MPC strategies thinning of bay is carried out early before switching to thinning of redwood. These species are reduced to a threshold density that OCT has identified as sufficiently low to suppress pathogen spread, and promote tanoak restoration. This type of insight about optimal densities of different species in mixed stands over the course of an epidemic could provide actionable advice for foresters.

The complexity of the optimal strategies found comes in part from balancing multiple costs and benefits: tanoak retention, biodiversity conservation, control expenditure and surveillance costs. The valuation of the cultural and ecological benefits against more direct economic costs is a difficult decision that must be taken by forest managers. These decisions must be made in the context of the local area, as well as other forest management goals such as fire reduction and timber production [18]. Decisions about management goals however, either locally or through larger scale regulation, can lead to conflicts that may impact the effectiveness of control [45]. The value of tanoak retention must be balanced with the wider impacts of the control on the forest.

As well as valuation, the formulation of the different cost functions is important. Here, we used a metric for biodiversity conservation which was integrated over time and so ensures biodiversity is conserved at all times. This captures the importance of continued biodiversity for wildlife habitats, but also avoids introducing edge effects, for example thinning very late in the epidemic to meet a biodiversity target. Both the biodiversity and tanoak objectives introduce a dependence on the chosen time horizon of 100 years, but the tanoak objective only depends on the amount of tanoak at the final time. This final time dependence is appropriate for a restoration type management goal, such as ensuring a resource is available in the future. There is still however, a flexibility in the form of the objective function chosen, and the precise choice of objective does impact disease control [46]. Extending the final time out to 160 years however, results in functional extinction of tanoak in the open-loop case (i.e. no significant ecosystem function), but retention of overstorey tanoak when using MPC (S3 Text).

We have shown that by repeated surveillance and re-optimisation of control, the damaging effects of the worst-case scenarios of pathogen spread can be limited. As found by Cobb *et al*. [18], disease control is only effective when there is long-term commitment to management projects. Effort put into this long-term surveillance has to be cost-effective though. With imperfect surveillance introducing observational uncertainty, an optimal balance between survey costs and epidemic control was found. Other modelling studies of SOD have also found that resource constraints lead to a trade off between detection and control [3, 47]. However, our analysis does not incorporate the risk of disease re-emergence. Our model is deterministic, so also does not capture stochastic re-introductions. Furthermore, the state of the epidemic in the wider region will impact the risk of re-emergence through potentially increasing inoculum pressure, for example from the advancing wavefront of other epidemics in the local vicinity. Forest managers must take into account these other factors in determining optimal levels of surveillance as well as in designing controls, but regardless we have shown that vigilance to disease progression is important. Alongside this vigilance must be a willingness to adapt control measures, re-optimising control to suit the current state of the epidemic and changing local management goals.

In testing robustness we showed that the MPC framework is able to mitigate the effects of the most damaging epidemics, improving management performance in the worst-case scenarios. Maccleery [48] states that a major barrier to US Forest Service management is the opposition to adaptive management, in which ongoing monitoring is used to update management advice. This is seen as too ‘experimental’ and increases short-term risk, but imposing fixed interventions means strategies cannot be adapted based on what is seen ‘on the ground’. Here we have shown a clear benefit to the ongoing surveillance and re-optimisation of control, with MPC as a possible formal framework for adapting strategies. The open-loop framework however, still significantly slowed tanoak decline, and slowing pathogen spread is still a useful goal allowing time to prepare for ecosystem impacts [49].

In this paper we have optimised strategies for slowing, or even halting, pathogen-induced decline of tanoak in mixed species forest stands. OCT was used to find optimal time-dependent deployment of thinning, roguing, and protectant control resources. The strategies found are broadly consistent with current expert advice: focussing on thinning of bay laurel and roguing of infected trees. However, the strategies we found show significant time dependence. Continued surveillance and re-optimisation of the control strategy by using the MPC framework improves control performance. MPC leads to robust strategies that can effectively respond to unanticipated disease dynamics and system uncertainties, and so manage SOD to protect valuable tanoak trees whilst also conserving biodiversity. Our future work will extend these frameworks to more complex spatial strategies, also designed to protect valuable forest resources.

## Methods

### Simulation model

The spatial simulation model used is an implementation of the model described by Cobb *et al*. [16]. A system of ordinary differential equations track the stem density (i.e. the number of tree stems per unit area) of redwood, susceptible and infected bay laurel, and susceptible and infected tanoak in four age classes. The model is spatial with hosts positioned on a 20 by 20 grid of square cells, each of area 500 m^2^. Recruitment and age transitions occur within a single cell, with density-dependence in the recruitment rates based on the available space in the cell. Infection dynamics are the only interaction between cells. In the original model [16], infected cells exerted an infectious pressure on susceptible hosts within the same cell and in the 4 adjacent cells. Infectious spores are distributed such that 50 % land within the same cell, and the other 50 % are distributed across the adjacent cells. In our model we use a more realistic dispersal kernel, with 50 % of spores landing within the same cell, and the other 50 % distributed according to an exponential kernel with a scale parameter of 10 m. The dispersal scale is chosen to be representative of the distance of local splash dispersal [50], and leads to dynamics with invasion rates consistent with field studies [13]. The kernel is normalised so that total spore deposition across all cells is 100 %. With this new spore deposition kernel, the infection rates are rescaled to match the time scales of invasion found by Cobb *et al* [16] (S1 Text).

Epidemics are seeded in the centre of a 20 by 20 grid (any one of the four central cells). Infection starts in the bay population and the smallest tanoak age class, with 50 % of these hosts starting infected. The forest host composition is taken from that used by Cobb *et al*. [16] for mixed species forest stands. This corresponds to 40 % tanoak, 16 % bay, and 44 % redwood. The initial amount of empty space is found by initialising the model in dynamic equilibrium (S1 Text). The state variables track stem density in arbitrary units. The model is parameterised from the study plots in Cobb *et al*. [16], which have an average tanoak density of 561 ha^-1^. The stem densities from the model can therefore be scaled such that the initial stem density is 1,400 ha^-1^, since tanoak makes up 40 % of the forest composition.

### Approximate model and fitting

To allow optimisation of a time-dependent control strategy, the simulation model is approximated by a simpler, non-spatial model. To ensure that we can lift demographic parameters directly from the simulation model however, we only change the spatial structure of the model (S2 Text). The approximate model therefore assumes that all hosts in the forest stand are well-mixed. The infection rates are fitted using a least squares approach, matching the disease progress curves to those from the simulation model (S2 Text).

### Control optimisation

Three classes of control are optimised: roguing, thinning and protectant methods. Roguing methods are based on finding and removing infected hosts, whereas thinning methods remove hosts regardless of infection status. Removal of hosts, either through thinning or roguing, is the only control that has been effective at the landscape scale [51]. Protectant methods apply chemicals to uninfected trees to reduce their susceptibility to the disease. For SOD the main protectants used are phosphonates, that are approved for use on oak and tanoak species and slow infection for up to two years [52]. Roguing controls can be applied separately to infected small tanoak, large tanoak and bay laurel. The hosts are removed and do not resprout, consistent with application of a herbicide to the stump as is often recommended [24]. Thinning removes hosts of all infection statuses, and can be applied separately to small tanoak, large tanoak, bay and redwood. Protection can only be applied to small and large tanoaks, and only to susceptible hosts. These hosts are moved into a new protected class with reduced susceptibility (S1 Text). Overall this gives 9 time-dependent control functions (*f*_*i*_ (*t*)).

The time-dependent controls are chosen to maximise an objective incorporating tanoak retention and biodiversity conservation. Since the focus of the control is to ensure overstorey tanoak is retained in the forest in the future, we treat the tanoak term as a terminal objective function, maximising the density of healthy overstorey tanoak after *T* = 100 years. The Shannon index is used as a metric for biodiversity [53], integrated over time to ensure ecosystem services are maintained throughout the epidemic. The total objective to be maximised is therefore given by:

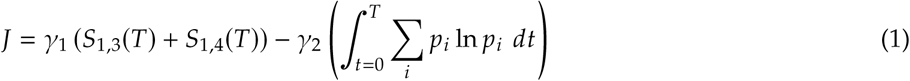

where *S*_1,3_ and *S*_1,4_ are the densities of healthy tanoak in the third and fourth age classes respectively, and *γ*_1_ and *γ*_2_ are the weights associated with the tanoak retention and biodiversity conservation objectives respectively. The weights are set such that in the disease free case, the contribution of the tanoak retention and biodiversity terms to the overall objective function are equal to 1 and 0.25 respectively. The sum in the second term is over host species, where *p*_*i*_ is the proportion of hosts in species *i*. The objective is maximised by optimising the time-dependent control functions using BOCOP v2.0.5 [54], subject to a budget constraint limiting the expenditure on control per unit time (S1 Text).

### Lifting and control frameworks

Strategies optimised using the approximate model are lifted to the simulation model. The control functions from the optimisation in the approximate model are applied to the simulation model, although distributed evenly across the landscape to give a non-spatial control strategy. The functions are applied ensuring adherence to the budget constraint (S3 Text). In the open-loop framework the control strategy is lifted to the simulation model and applied for the full 100 years. In the MPC framework every 20 years the approximate model is reset to match the current state of the simulation model. The initial conditions in the approximate mode are set to match the average density of each host across the landscape in the simulation model. Control is then re-optimised using the approximate model and lifted to the simulation for the next 20 years.

### Parameter uncertainty

For simplicity we assume uncertainty is only present in the infection rates. A truncated normal distribution is used to perturb the infection rates and ensure they remain positive, with a standard deviation of 40 % of the parameter value. An ensemble of 250 simulations is used to fit the approximate model to the average behaviour across the ensemble, again using a least squares approach. The open-loop and MPC frameworks are then tested using the fitted approximate model and new draws from the parameter distribution for the simulation model.

### Observational uncertainty

At each MPC update time a proportion of the forest stand is sampled. The measured state is then used as the initial condition in the approximate model for re-optimisation of the control. For the sampling, each cell in the simulation model is split into 500 1 m^2^ discrete units. Surveillance at update times is then carried out by observing a fixed number of units in each cell across the landscape, without replacement. The infection status of tanoak and bay hosts is determined randomly, with probabilities matching the proportion of that host that is infected.

## Supporting information

**S1 Figure. Host dynamics under MPC with low biodiversity benefit.** The optimal control strategy using the MPC framework when there is no biodiversity benefit removes all bay and redwood from the forest.

**S1 Text. Simulation model details.** Full details and parameterisation of the simulation model, and adaptations made to the original implementation. The control methods and budget constraint are also explained.

**S2 Text. Approximate model and fitting details.** The formulation of the non-spatial approximate model and the methods used to fit it to simulation data.

**S3 Text. Control frameworks and optimisation results.** Description of the open-loop and MPC frameworks and methods for ensuring adherence to the budget constraint. We also here show the effect of varying the frequency of update times, extending the time horizon, and removing protectant application.

**S4 Text. Parameter uncertainty.** Here we analyse the control strategies and what drives changes in performance for the four parameter uncertainty scenarios from Fig 5.

## Additional Information

## Acknowledgements

We thank Richard Cobb for helpful discussions and provided code for the original model implementation.

## Data Accessibility

All code and generated data are available at https://github.com/ehbussell/MixedStand.

## Authors’ Contributions

E.H.B and N.J.C. designed the study, E.H.B. conducted the analysis and wrote the initial draft of the manuscript. Both authors contributed to data interpretation, manuscript editing and discussion.

