## Supplementary material for "Ongoing surveillance protects tanoak whilst conserving biodiversity: applying optimal control theory to a spatial simulation model of sudden oak death": S1 Figure

### Supporting Information S1 Figure

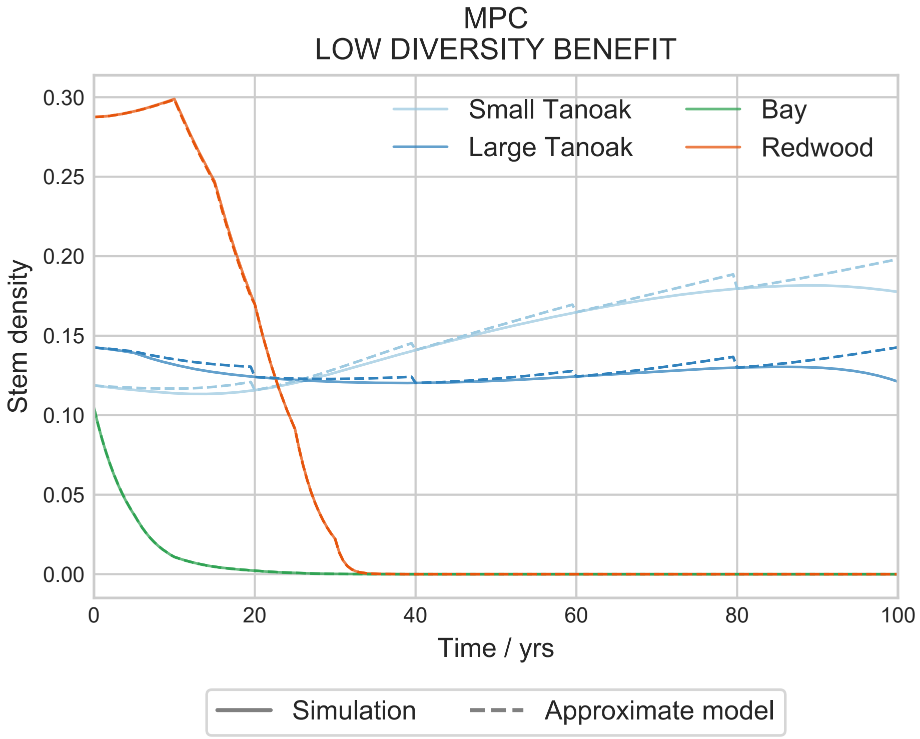


**Fig S1. Host dynamics under MPC with low biodiversity benefit.** The optimal control strategy using the MPC framework when there is no biodiversity benefit removes all bay and redwood from the forest.
