## Supplementary material for "Ongoing surveillance protects tanoak whilst conserving biodiversity: applying optimal control theory to a spatial simulation model of sudden oak death": S1 Text

### Supporting Information S1 Text

E.H. Bussell & N.J. Cunniffe

#### S1 Simulation model details

##### S1.1 Model description

The simulation model used in the main text is adapted from the sudden oak death (SOD) model in Cobb et al. [2012]. We here describe initially the model as implemented in Cobb et al. [2012], and then the modifications made in our implementation. As described in the main text, the model tracks the stem density dynamics of three different host species or groups: redwood, bay laurel and tanoak. The redwood class comprises of all epidemiologically inactive hosts, which for the stands considered is predominantly coast redwood. The tanoak class is divided into 4 separate age groups, in order to capture the effects of disease on the older trees, with the two oldest groups corresponding to the overstorey tanoak. As differing effects on older trees are less important for the other hosts because bay is not killed by the disease, and to reduce model complexity, the other host groups are not divided into age classes. The model tracks natural host demography, with natural mortality and seed recruitment rates allowed to differ for each host class. Recruitment depends on the amount of empty space available for seedling establishment, with each tanoak age class weighted to occupy differing amounts of space per stem. Over time tanoak hosts progress through the age classes. See Figure 1A in the main text for an overview of the different classes and possible transitions.

The model is spatially-explicit, with hosts positioned on a $20 \times 20$ grid in square cells each of area $500\,\mathrm{m}^2$ and 400 cells in total. Recruitment and age transitions occur at rates based entirely on host density within a single cell, with density-dependence in the recruitment rates based on the available space in the cell. Infection dynamics are therefore the only interaction between cells. Infected hosts exert infectious pressure on susceptible hosts within the same cell and in the 4 adjacent cells. Infectious spores are distributed such that 50 % land within the same cell, and the other 50 % are distributed across the adjacent cells. Bay and all age classes of tanoak are susceptible, and are infectious once infected without a latent period.

The model is formulated as a system of ODEs resulting in 11 differential equations per cell. We use $S$ to indicate healthy hosts, $I$ to indicate infected hosts, and subscripts to indicate species (1: tanoak, 2: bay, 3: redwood), age class (1–4 where applicable), and cell location ($x$), following the notation used in Cobb et al.

[2012] and as shown in Figure 1A in the main text. The resulting equations for cell $x$ and age class $i$ are:

$$\begin{aligned}\dot{S}_{1,i,x} &= \delta_{1,i}\left[B_{1,x}E_x + r\alpha_{1,i}I_{1,i,x}\right] - d_{1,i}S_{1,i,x} - \Lambda_{1,i,x}S_{1,i,x} + \mu_1 I_{1,i,x} \\ &\quad + \left[1-\delta_{1,i}\right]a_{i-1}S_{1,i-1,x} - \left[1-\delta_{4,i}\right]a_i S_{1,i,x}\end{aligned} \tag{1a}$$

$$\begin{aligned}\dot{I}_{1,i,x} &= -\alpha_{1,i}I_{1,i,x} - d_{1,i}I_{1,i,x} + \Lambda_{1,i,x}S_{1,i,x} - \mu_1 I_{1,i,x} \\ &\quad + \left[1-\delta_{1,i}\right]a_{i-1}I_{1,i-1,x} - \left[1-\delta_{4,i}\right]a_i I_{1,i,x}\end{aligned} \tag{1b}$$

$$\dot{S}_{2,x} = b_2\left(S_{2,x} + I_{2,x}\right)E_x - d_2 S_{2,x} - \Lambda_{2,x}S_{2,x} + \mu_2 I_{2,x} \tag{1c}$$

$$\dot{I}_{2,x} = -d_2 I_{2,x} + \Lambda_{2,x}S_{2,x} - \mu_2 I_{2,x} \tag{1d}$$

$$\dot{S}_{3,x} = b_3 S_{3,x}E_x - d_3 S_{3,x} \tag{1e}$$

where tanoak dynamics are given by equations 1a and 1b, bay dynamics by 1c and 1d, and redwood dynamics by 1e (recall that all hosts in this class cannot become infected). All parameter meanings and symbols are given in Table 1. The delta function at the start of equation 1a, $\delta_{1,i}$, is equal to one for the smallest age class, and zero otherwise. This ensures that recruitment is always to the smallest age class.

The recruitment rates are density dependent as they depend on the space available for colonisation in each cell, $E_x$. The empty space in cell $x$ is given by:

$$E_x = 1 - W_1 \sum_{i=1}^{4} w_{1,i}\left(S_{1,i,x} + I_{1,i,x}\right) - W_2\left(S_{2,x} + I_{2,x}\right) - W_3 S_{3,x}\ . \tag{2}$$

The species suppression weights $W_j$ give the relative area colonised, and hence unavailable for seedling recruitment, by each species per capita, which are assumed to all be equal to 1. The tanoak suppression weights $w_{1,i}$ give different relative space occupation for each tanoak age class. These suppression weights capture actual space occupied, as well as seedling suppression by other means, for example by blocking sunlight. The tanoak recruitment rate is made up of each individual recruitment from each age class, but all seedlings enter at the smallest age class. The tanoak recruitment rate, $B_{1,x}$, in equation 1a is given by:

$$B_{1,x} = \sum_{i=1}^{4} b_{1,i}\left(S_{1,i,x} + I_{1,i,x}\right) \tag{3}$$

so that recruitment is from all age classes, with seed production rates that are independent of infection status.

Finally we describe the force of infection terms $\Lambda$ in equations 1:

$$\Lambda_{1,i,x} = f_0\left[\beta_{1,i}\sum_{j=1}^{4} I_{1,j,x} + \beta_{12}I_{2,x}\right] + f_1 \sum_{y \in N(x)}\left[\beta_{1,i}\sum_{j=1}^{4} I_{1,j,y} + \beta_{12}I_{2,y}\right] \tag{4a}$$

$$\Lambda_{2,x} = f_0\left[\beta_{21}\sum_{j=1}^{4} I_{1,j,x} + \beta_{2}I_{2,x}\right] + f_1 \sum_{y \in N(x)}\left[\beta_{21}\sum_{j=1}^{4} I_{1,j,y} + \beta_{2}I_{2,y}\right] \tag{4b}$$

where $\beta_{12}$ is the rate of infection from bay to tanoak, $\beta_{21}$ from tanoak to bay, and $\beta_2$ within bay. The infection

Table 1: Parameter values used in Cobb et al. [2012]. Parameter values marked with an asterisk are set in model initialisation to impose dynamic equilibrium. See Section S1.2 for a full description.

| **Parameter** | | **Symbol** | **Default Value** |
|---|---|---|---|
| Infection rate | tanoak to tanoak (1 cm to 2 cm d.b.h.) | $\beta_{1,1}$ | 0.33 $\text{year}^{-1}$ |
| | tanoak to tanoak (2 cm to 10 cm d.b.h.) | $\beta_{1,2}$ | 0.32 $\text{year}^{-1}$ |
| | tanoak to tanoak (10 cm to 30 cm d.b.h.) | $\beta_{1,3}$ | 0.30 $\text{year}^{-1}$ |
| | tanoak to tanoak (>30 cm d.b.h.) | $\beta_{1,4}$ | 0.24 $\text{year}^{-1}$ |
| | bay to tanoak | $\beta_{12}$ | 1.46 $\text{year}^{-1}$ |
| | bay to bay | $\beta_{2}$ | 1.33 $\text{year}^{-1}$ |
| | tanoak to bay | $\beta_{21}$ | 0.30 $\text{year}^{-1}$ |
| Natural mortality rate | tanoak (1 cm to 2 cm d.b.h.) | $d_{1,1}$ | 0.006 $\text{year}^{-1}$ |
| | tanoak (2 cm to 10 cm d.b.h.) | $d_{1,2}$ | 0.003 $\text{year}^{-1}$ |
| | tanoak (10 cm to 30 cm d.b.h.) | $d_{1,3}$ | 0.001 $\text{year}^{-1}$ |
| | tanoak (>30 cm d.b.h.) | $d_{1,4}$ | 0.032 $\text{year}^{-1}$ |
| | bay | $d_{2}$ | 0.02 $\text{year}^{-1}$ |
| | redwood | $d_{3}$ | 0.02 $\text{year}^{-1}$ |
| Recruitment rate | tanoak total cell $x$ | $B_{1,x}$ | Equation 3 |
| | tanoak (1 cm to 2 cm d.b.h.) | $b_{1,1}$ | 0.0 $\text{year}^{-1}$ |
| | tanoak (2 cm to 10 cm d.b.h.) | $b_{1,2}$ | 0.007 $\text{year}^{-1}$ |
| | tanoak (10 cm to 30 cm d.b.h.) | $b_{1,3}$ | 0.02 $\text{year}^{-1}$ |
| | tanoak (>30 cm d.b.h.) | $b_{1,4}$ | 0.073 $\text{year}^{-1}$ |
| | bay | $b_{2}$ | * |
| | redwood | $b_{3}$ | * |
| Disease induced mortality rate | tanoak (1 cm to 2 cm d.b.h.) | $\alpha_{1,1}$ | 0.019 $\text{year}^{-1}$ |
| | tanoak (2 cm to 10 cm d.b.h.) | $\alpha_{1,2}$ | 0.022 $\text{year}^{-1}$ |
| | tanoak (10 cm to 30 cm d.b.h.) | $\alpha_{1,3}$ | 0.035 $\text{year}^{-1}$ |
| | tanoak (>30 cm d.b.h.) | $\alpha_{1,4}$ | 0.14 $\text{year}^{-1}$ |
| Tanoak age transition rate | (1 cm to 2 cm d.b.h.) to (2 cm to 10 cm d.b.h.) | $a_{1}$ | 0.142 $\text{year}^{-1}$ |
| | (2 cm to 10 cm d.b.h.) to (10 cm to 30 cm d.b.h.) | $a_{2}$ | 0.2 $\text{year}^{-1}$ |
| | (10 cm to 30 cm d.b.h.) to (>30 cm d.b.h.) | $a_{3}$ | 0.05 $\text{year}^{-1}$ |
| Recovery rate | tanoak | $\mu_{1}$ | 0.01 $\text{year}^{-1}$ |
| | bay | $\mu_{2}$ | 0.1 $\text{year}^{-1}$ |
| Recruitment suppression weight | species $i$ | $W_{i}$ | 1 |
| | tanoak age class $i$ | $w_{1,i}$ | * |
| Resprouting probability | tanoak | $r$ | 0.5 |
| Spore proportion | within cell | $f_{0}$ | 0.5 |
| | between cell | $f_{1}$ | 0.125 |
| Force of infection | tanoak age class $i$, cell $x$ | $\Lambda_{1,i,x}$ | Equation 4a |
| | bay, cell $x$ | $\Lambda_{2,x}$ | Equation 4b |

rate within tanoak is given by $\beta_{1,i}$, meaning each age class has a different susceptibility to infection from other tanoaks. Overall however, tanoak age classes do not vary in susceptibility to infection from bay, nor in the rate of infecting bay. Hence, $\beta_{12}$ and $\beta_{21}$ do not depend on age class. The parameters $f_0$ and $f_1$ give the proportion of spores deposited within and between cells respectively, where the sum over cells $N(x)$ is over the four cells adjacent to $x$.

#### S1.2 Spore deposition and reparameterisation

In our implementation we use a more realistic spore deposition pattern than in the original model, by introducing an exponential dispersal kernel. The same proportion of spores (50 %) are deposited within the source cell as used in Cobb et al. [2012], corresponding to $f_0 = 0.5$. The other 50 % are distributed to cells other than the source cell according to an exponential kernel with a scale parameter of 10 m. The kernel is normalised so that total spore deposition across all cells is 100 %. The spore proportion between cells becomes proportional to:

$$\exp\left(-d_{ij}/\sigma\right) \tag{5}$$

where $d_{ij}$ is the distance from source cell to target cell, and $\sigma$ is the scale parameter. The choice of 10 m as a scale parameter is somewhat arbitrary, although consistent with distances of splash dispersal found for *P. ramorum* [Davidson et al., 2005] and equal to the mean dispersal distance used by Cobb et al. [2012].

The bay and redwood birth rates ($b_2$ and $b_3$) are fixed to give dynamic equilibrium, following the same process used by Cobb et al. [2012]. The initial amount of empty space ($E_x(0)$) and the initial tanoak age distribution are also found by imposing dynamic equilibrium on Equations 1, but here found analytically unlike in the original model. The change to the spore deposition kernel means the dynamics are now different from those reported by Cobb et al. [2012]. The time scales for invasion found by Cobb et al. [2012] are consistent with expectations [Davidson et al., 2005, McPherson et al., 2010], and so we rescale our implementation to give the same rate of invasion. We use the time at which the population of small tanoak increases above the large tanoak population to define the rate of invasion. All infection rates in our implementation are then scaled by the same factor so that relative rates are kept the same, but the rate of invasion matches that of the original Cobb model implementation (Figure 1). This ensures that the dynamics are correct, with a realistic kernel distribution, whilst matching the generally accepted spread rates for SOD. Our choice for defining the time scale is arbitrary, but since the results show that the best fit is very close to the original at all times, other choices would not give very different results.

Since we are reparameterising the model through the rescaling, we also take the opportunity to correct another unrealistic parameterisation in the orginal model. The tanoak suppression weights $w_{1,i}$ in the original model are chosen such that a quarter of space is taken up by each age class. For the age distribution used this meant the greatest suppression was unrealistically from the smallest age class. In our implementation, the weights are chosen to scale approximately with basal area. This ensures that younger age classes occupy less space than the older classes. The code for the original model also had the highest recruitment rate in the smallest age class, which we have corrected to be zero in our implementation following the parameters given in the paper [Cobb et al., 2012].

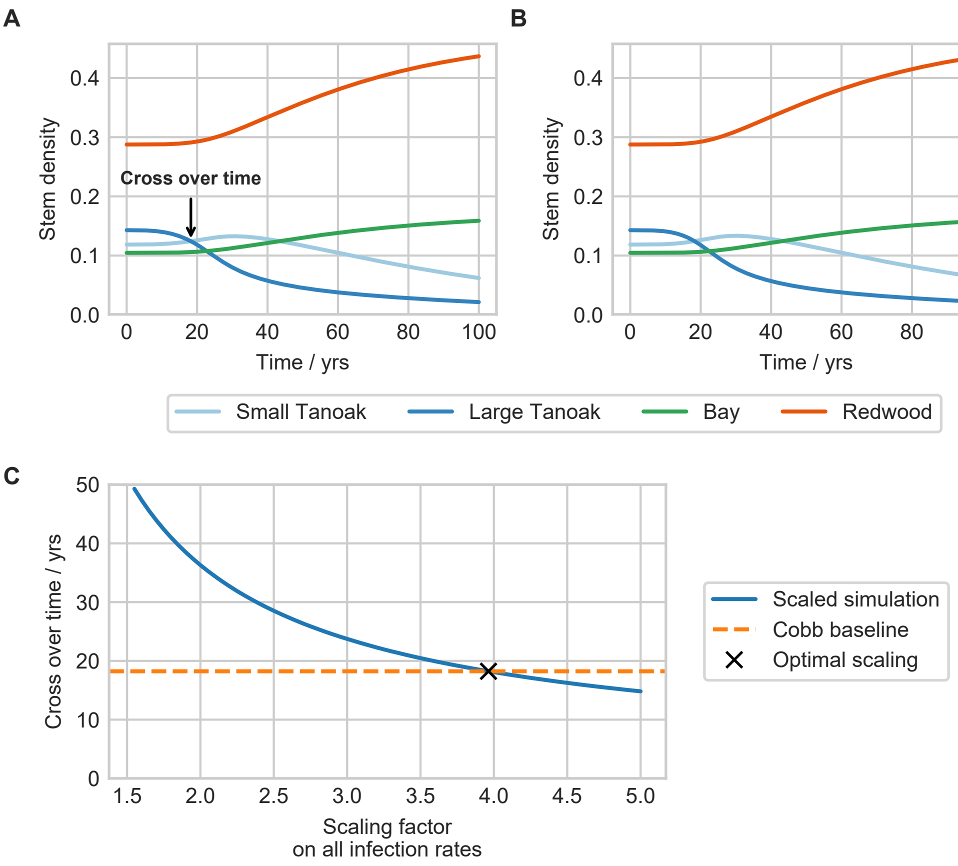


Figure 1: Effects of exponential spore deposition kernel, and matching time scales. A: the original dynamics in Cobb et al. [2012], with the cross over time labelled. The cross over time is the time until the large tanoak population falls below the small tanoak population, and is the metric we have chosen to measure the time scale of the epidemic. B: the dynamics using the exponential kernel, with the epidemic time scale matched to that in A. This is matched by scaling the infection rates, with the effect of this scaling factor on the cross over time shown in C. The optimal scaling factor is found to be 3.96 (3 s.f.).

#### S1.3 Control methods

As explained in the main text, we implement roguing, thinning and protectant controls in the model. Roguing controls can be applied separately to infected small tanoak, large tanoak and bay laurel. The hosts are removed and do not resprout, consistent with application of a herbicide to the stump as is often recommended [Swiecki and Bernhardt, 2013]. Thinning removes hosts of all infection statuses, and can be applied separately to small tanoak, large tanoak, bay and redwood. Protection can only be applied to small and large tanoaks, and only to susceptible hosts. These hosts are moved into new protected class with the same demographic dynamics (i.e. there is a protected class $P_{1,i}$ for each age class $i$ of tanoak). The protected classes have reduced susceptibility (by 25 %) but return to the susceptible class at a rate of 0.5 year$^{-1}$. This corresponds to an average time of 2 years before protection wanes. Table 2 summarises all the control methods and their effects.

The application of control is optimised subject to a budget constraint. We seek a time-varying control parameter $f_i(t)$ between zero and one, indicating the level of control $i$, that minimises the management objective function given in the main text. To model economic and logistic constraints we limit the total expenditure per unit time, where this is the product of the number of hosts controlled and the cost of that control method. The mathematical form of this constraint is given by:

$$\sum_i \left(f_i \eta_i X_i\right) c_i \leq B \tag{6}$$

where $X_i$ is the stem density of the controlled hosts. For example, for roguing of small tanoak $X_i$ would be $(I_{1,1} + I_{1,2})$. The term in brackets is therefore the rate of removal of hosts for each control. The cost of each control is given by $c_i$ and the maximum budget is given by $B$, with a default value of 16 arbitrary units. Whilst the costs are chosen somewhat arbitrarily because of a lack of data, the scales are informed by the results of Kovacs et al. [2011]. We include higher costs for roguing to capture the additional costs with identification and removal of unstable infected trees.

Table 2: Possible control methods implemented in the stand level model. There are three main groups of control: roguing, thinning and protecting, and these can be targeted at different host groups. Approximate costs of each control are taken from Kovacs et al. [2011], but roguing costs are increased to account for additional costs of identification and removal of unstable diseased trees.

| **Control** | **State changes** | **Rate** $\eta_I$ / year$^{-1}$ | **Cost** $c_i$ / a.u. |
|---|---|---|---|
| Rogue small tanoak | $I_{1,1\text{–}2} \rightarrow \emptyset$ | 0.25 | 3000 |
| Rogue large tanoak | $I_{1,3\text{–}4} \rightarrow \emptyset$ | 0.25 | 6000 |
| Rogue bay | $I_2 \rightarrow \emptyset$ | 0.25 | 6000 |
| Thin small tanoak | $\{S, I, P\}_{1,1\text{–}2} \rightarrow \emptyset$ | 1.0 | 250 |
| Thin large tanoak | $\{S, I, P\}_{1,3\text{–}4} \rightarrow \emptyset$ | 1.0 | 500 |
| Thin bay | $\{S, I\}_2 \rightarrow \emptyset$ | 1.0 | 500 |
| Thin redwood | $S_3 \rightarrow \emptyset$ | 1.0 | 500 |
| Protect small tanoak | $S_{1,1\text{–}2} \rightarrow P_{1,1\text{–}2}$ | 0.25 | 200 |
| Protect large tanoak | $S_{1,3\text{–}4} \rightarrow P_{1,3\text{–}4}$ | 0.25 | 200 |



©The Authors under the terms of the Creative Commons Attribution License `http://creativecommons.org/licenses/by/3.0/`, which permits unrestricted use, provided the original author and source are credited.
