## Supplementary material for "Ongoing surveillance protects tanoak whilst conserving biodiversity: applying optimal control theory to a spatial simulation model of sudden oak death": S2 Text

### Supporting Information S2 Text

E.H. Bussell & N.J. Cunniffe

#### S2 Approximate model and fitting details

##### S2.1 Approximate model

The approximate model used to optimise the control via OCT is non-spatial, but in all other respects is identical to the simulation model. The demographic parameters, such as age transition rates and birth rates, can be used directly from the simulation model since the demographic dynamics are the same. The infection rates in the non-spatial approximate model cannot be lifted from the simulation model, since the approximate model now assumes infection comes from all infected hosts in the stand rather than through the spore deposition kernel. The force of infection terms in the approximate model are therefore:

$$\tilde{\Lambda}_{1,i} = \tilde{\beta}_{1,i} \sum_{j=1}^{4} I_{1,j} + \tilde{\beta}_{12} I_2 \tag{1a}$$

$$\tilde{\Lambda}_2 = \tilde{\beta}_{21} \sum_{j=1}^{4} I_{1,j} + \tilde{\beta}_2 I_2 \tag{1b}$$

where $\tilde{\beta}$ indicate infection parameters that need to be fitted to the simulation model, and the dependence on cell ($x$) of the states has been dropped since the model is non-spatial.

##### S2.2 Fitting infection rates

The seven infection rates $\tilde{\beta}$ are the only parameters that must be fitted to the simulation model. We use the method of least squares to match the simulation and approximate models. To fit the parameters, the simulation model is used to run a single trajectory with no control interventions. For the fitting process we use the same initial conditions as described in the main text, with infection seeded in the centre of a 20 by 20 grid of cells. The disease progress curves of this simulation run are then used as the baseline for fitting the approximate model. For a trial set of $\tilde{\beta}$ parameters and an approximate model trajectory, we calculate the sum of squares as the sum of squared deviations between the simulation and approximate disease progress curves for each age class of tanoak, and for bay, at time points throughout the trajectory. Redwood is not included as it cannot be infected. The $\tilde{\beta}$ parameters are then optimised by minimising this total summed squared error (SSE). For a set of time points $t_i$, and where approximate model states are signified with a tilde, the equation for SSE is given by:

$$\text{SSE} = \sum_i \left[ \sum_{j=1}^{4} \left( I_{1,j}(t_i) - \tilde{I}_{1,j}(t_i) \right)^2 + \left( I_2(t_i) - \tilde{I}_2(t_i) \right)^2 \right] \tag{2}$$

where the dependence on cell in the simulation terms has been dropped to indicate an average over all cells in the landscape, for example:

$$I_{1,j}(t) = \frac{\sum_x I_{1,j,x}(t)}{N_{\text{cells}}} \ . \tag{3}$$

An average is used so that the approximate model tracks stem density in the same units as the simulation model.

In the simulation model, infectious pressure is dominated by sporulation from bay laurel. This makes estimation of all 'within-tanoak' infection rates ($\beta_{1,i}$) difficult, as from the simulation data they are individually unidentifiable. We therefore use a two stage fitting process. For the first stage, all infection rates in the simulation model related to bay ($\beta_2$, $\beta_{12}$, and $\beta_{21}$) are set to zero. This makes bay epidemiologically inactive, but maintains the same demographic dynamics. The simulation model is run using these parameters, and the SSE is minimised to find the within tanoak-infection rates.

In the second fitting stage all infection rates are fitted, with bay epidemiologically active again. The within-tanoak rates relative to $\beta_{1,1}$ from the first stage are used as a constraint to ensure the identifiability of these rates. This means a single within-tanoak rate is fitted in stage 2, with all other within-tanoak rates fixed relative to this using the results from stage 1. This stage also fits the bay infection rate, and the cross-species infection rates.

As can be seen in Figure 1B–1C in the main text, despite lacking any spatial component, the approximate model can very closely capture the uncontrolled dynamics of the simulation model. However, the approximate model should also fit as accurately as possible when control strategies are introduced. In Figure 1, the fit of the approximate model is tested under constant control strategies using fixed control rates. It is clear that roguing at the same rate is more effective in the approximate model. This is because of the difference in mixing between the approximate and simulation models. The effect is small for thinning and protecting, but the same level of roguing in the approximate and simulation models gives very different dynamics.

##### S2.2.1 Empirical parameterisation of control

Roguing is less effective in the simulation model because of an imposed spatial structure in the non-spatial control strategy. Roguing in the simulation model removes infected hosts from the core and edge of the spreading epidemic. Removal of hosts from the core has little effect on the rate of epidemic spread, since they are not near the wavefront. In the non-spatial model however, all hosts are well-mixed, so removal of infected hosts has a larger effect.

As a simple correction for this difference in roguing effectiveness, we investigate a simple scaling of the roguing rate in the approximate model. To test the plausibility of a single scaling rate for all approximate roguing controls, both models are run with constant roguing strategies. Roguing of small tanoak, large tanoak and bay are all set to occur at the same rate for the whole simulation, and this rate is varied between simulations. In the approximate model, the control rate is scaled by a single parameter which is also varied, and we analyse the difference in the final number of healthy large tanoak after 100 years between the simulation and approximation (Figure 2). To minimise the deviation across a range of control rates, the value of the scaling parameter is optimised. We optimise the deviation in the number of large healthy tanoak, since this is the primary objective of the control and must therefore be captured as accurately as possible. The optimisation minimised the sum of squared errors (SSE) over the range of control rates.

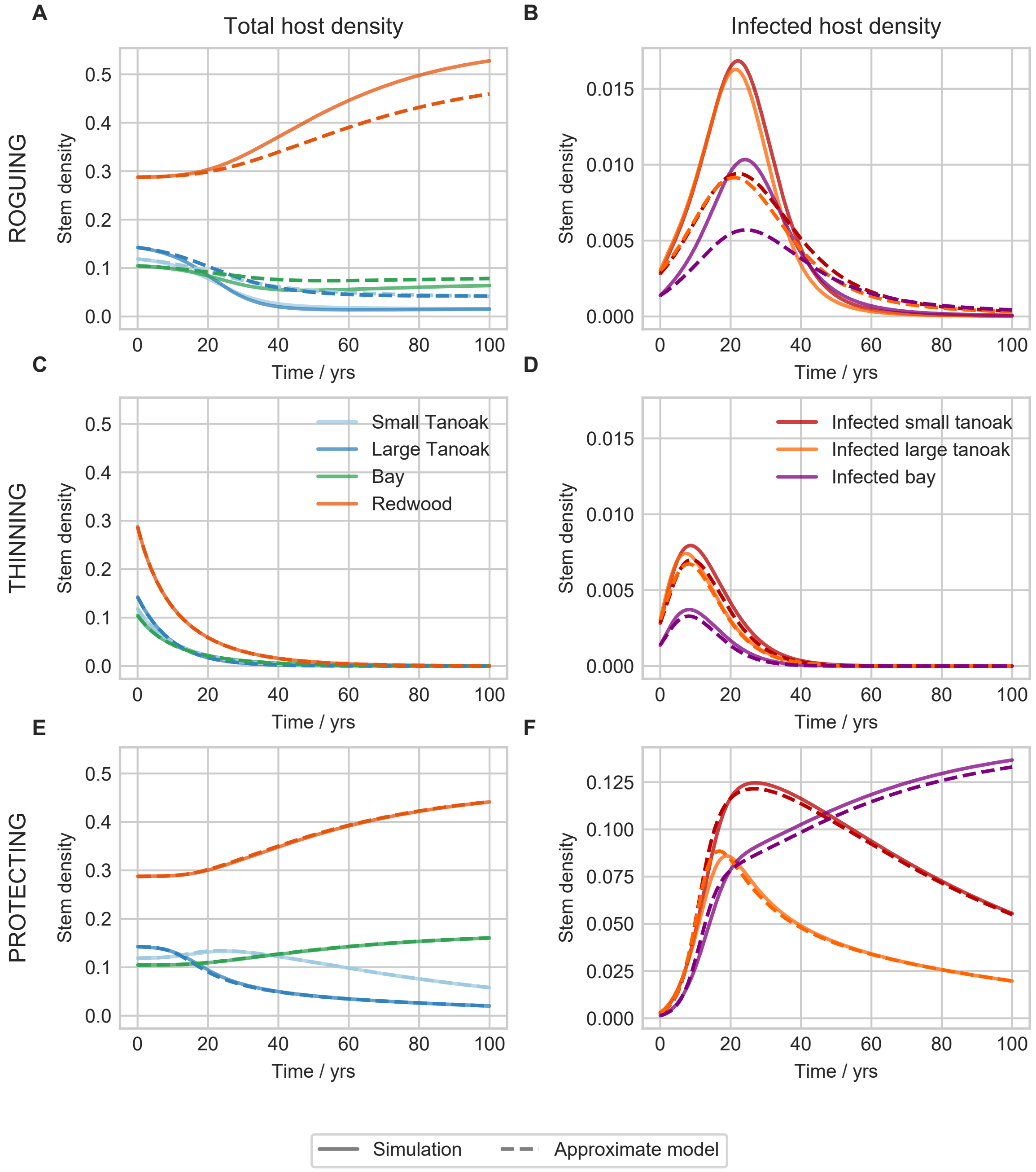


Figure 1: Testing of approximate model under constant control strategies. The rows from top to bottom show dynamics under constant roguing, thinning, and protecting strategies. The left plots show overall host dynamics, and the right plots show the infected host dynamics. The roguing and protecting strategies control at the maximum rates from Table 1 in S1 Text, whereas the thinning strategy controls at 10 % of the maximum rate. The approximate model fits well under constant thinning and protecting strategies, but less well under a constant roguing strategy.

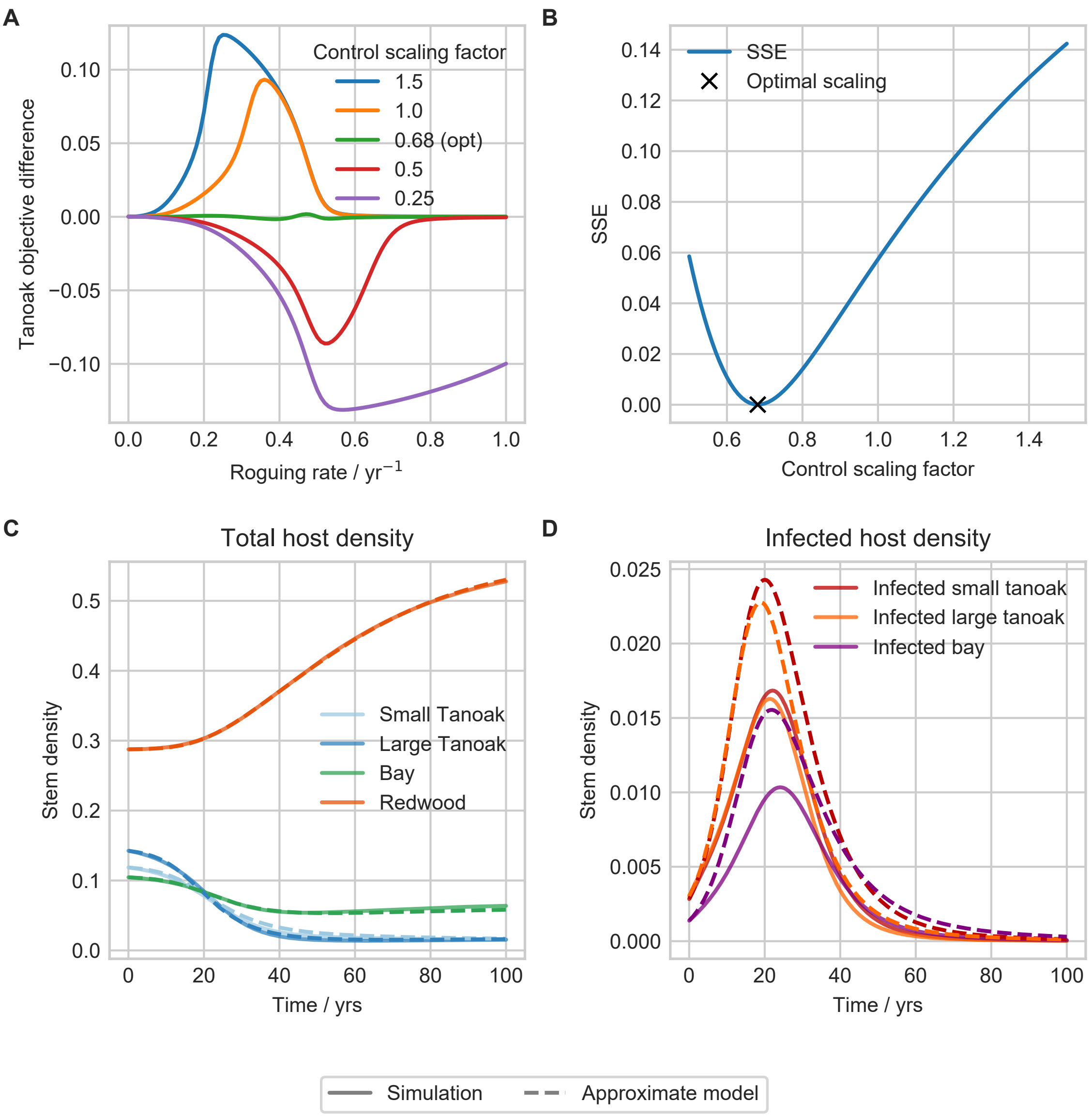


Figure 2: Empirical scaling of approximate model control rates to match simulation output. A: the difference in the final number of large healthy tanoak as a function of the constant roguing rate, for a number of control rate scaling factors. The optimal value minimises the sum of squared errors (SSE) over all rates, as shown in B, and is found to be 0.682 (3 s.f.). C and D show the dynamics with the newly scaled approximate control rates, under a constant roguing strategy. The approximate model now fits the overall host dynamics well, but to do this slightly overestimates the level of infection.

The results in Figure 2 show that a single scaling factor can largely eliminate the deviation under constant roguing strategies. We do not expect this scaling to ensure that the approximate model is always closely aligned to the simulation model, particularly once control strategies are time-varying. The approximate model simply cannot capture the heterogeneities in host mixing. This scaling does however go some way to ensuring that control strategies from the approximate model perform well on the simulation model. Figure 3 shows the open-loop and MPC control strategies and host dynamics when the roguing rate is not rescaled. Whilst OCT finds similar strategies, the error in the approximate model under open-loop is larger than when the roguing rate is scaled.

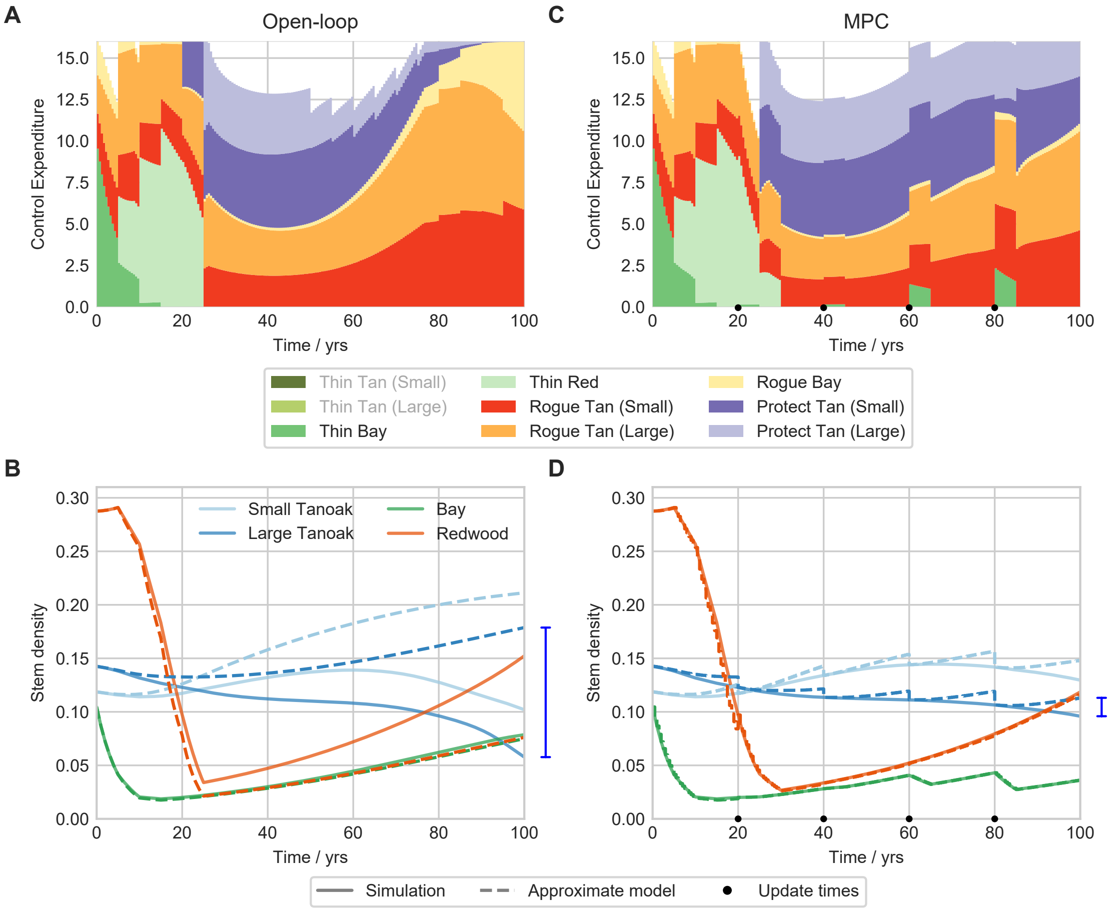


Figure 3: Control strategies and host dynamics under open-loop (A and B) and MPC (C and D) when the roguing rate is not scaled. Similar strategies are found but the error in the approximate model (blue bar) is much larger under open-loop than with the rescaling (compare with Figure 2 in the main text).

©The Authors under the terms of the Creative Commons Attribution License http://creativecommons.org/licenses/by/3.0/, which permits unrestricted use, provided the original author and source are credited.
