## Supplementary material for "Ongoing surveillance protects tanoak whilst conserving biodiversity: applying optimal control theory to a spatial simulation model of sudden oak death": S3 Text

### Supporting Information S3 Text

E.H. Bussell & N.J. Cunniffe

### S3 Control frameworks and optimisation results

#### S3.1 Control frameworks

The optimisation results from the approximate model are applied to the simulation model using the open-loop and model predictive control (MPC) frameworks described in Bussell et al. [2019]. In the open-loop framework, the control inputs ($f_i$) from the approximate model are applied directly to the simulation model for the whole simulation time. The budget constraint described above is a mixed constraint that couples the control inputs ($f_i$) with the state of the system ($X_i$). When the control inputs are lifted to the simulation model, there is no guarantee of the states being exactly the same, and so the expenditure by the control will not be the same. This can mean that direct lifting of the control inputs will lead to the budget being exceeded. When the budget is exceeded, the control inputs are multiplied by a time dependeent factor to reduce the overall expenditure to meet the budget constraint. This avoids imposing a priority amongst control methods which would have to be chosen arbitrarily. In mathematical terms, the control inputs from the approximate model depend on the state in the simulation ($X_i$) in the following way:

$$f_i(t) = f_i(t) \frac{B}{\sum_j \left( f_j(t) \eta_j X_j(t) c_j \right)} \ . \tag{1}$$

This is only the case when the budget is exceeded, otherwise the control inputs $f_i$ are used directly. We do not correct for under-allocation of resources, i.e. control inputs not spending the entire budget, since this could lead to extra control not accounted for in the approximate model. In practice correcting for this would lead to extra resources allocated to thinning, and hence removal of more healthy trees than is necessary.

In the MPC framework, at regular update times (every 20 years in the main text) the approximate model is reset to match the average stem density of each host across the landscape in the spatial simulation. The control is then re-optimised using these new initial conditions, and lifted back to the simulation model until the next update time. The same adjustment to the control inputs is made if the budget is exceeded. The MPC framework is described in Algorithm 1.

#### S3.2 Effect of update time frequency on MPC performance

Since the updates in MPC improve control, an important question is how often to update the approximate model. Figure 1A shows the effect of changing the update period on the objective. We can see that as updates are made more frequent, control performance generally improves. This is because the approximate model can more closely match the simulation and hence appropriate control decisions can be made. There is however, a

**Algorithm 1** MPC framework. MPC re-optimises the control at the update times.

1. Set initial conditions for simulation model
2. Fit approximate model to simulation data
3. Initialise approximate model at current simulator state (average host density)
4. Optimise control on approximate model
5. Lift control to simulation model and simulate forward
6. At next update time go to step 3

dip in performance at update periods of around 50 years. This is due to the precise timing of the updates. The late disease re-emergence occurs at around 80 years. Update periods of around 50 years will not update close to this outbreak, and so the MPC framework cannot respond to that unexpected increase in infection. This results in ineffective control.

Figures 1(b) and (c) show the MPC control for update periods of 5 and 100 years respectively. The low frequency update corresponds to open-loop control. We can see that the main difference as update frequency increases, is additional continued thinning of bay. This results in less roguing being required later in the epidemic.

#### S3.3 Extending the time horizon

The 100 year time horizon was chosen to be long enough to capture tanoak decline, and show the differences between the open-loop and MPC frameworks. We here extend the time horizon by 3 MPC update periods to 160 years to verify that the difference between the frameworks is robust in the longer term. Figure 2 shows the control strategies and host dynamics for both open-loop and MPC over this longer time horizon. The MPC framework does show tanoak decline from late re-emergence, but this is delayed compared to open-loop. After 160 years, tanoak is largely extinct under open-loop and tanoak numbers are still decreasing, but the MPC framework retains a stable population of large, healthy tanoak which is just beginning to increase in size.

#### S3.4 Efficacy of protectant methods

In the main text we state that the protectant methods are unlikely to be applied in practice, since they have a very small effect on the overall objective. We here verify this by running the MPC strategy with and without the protectant methods, as shown in Figure 3. The protectant application marginally increases the objective function, but the effect is negligible.

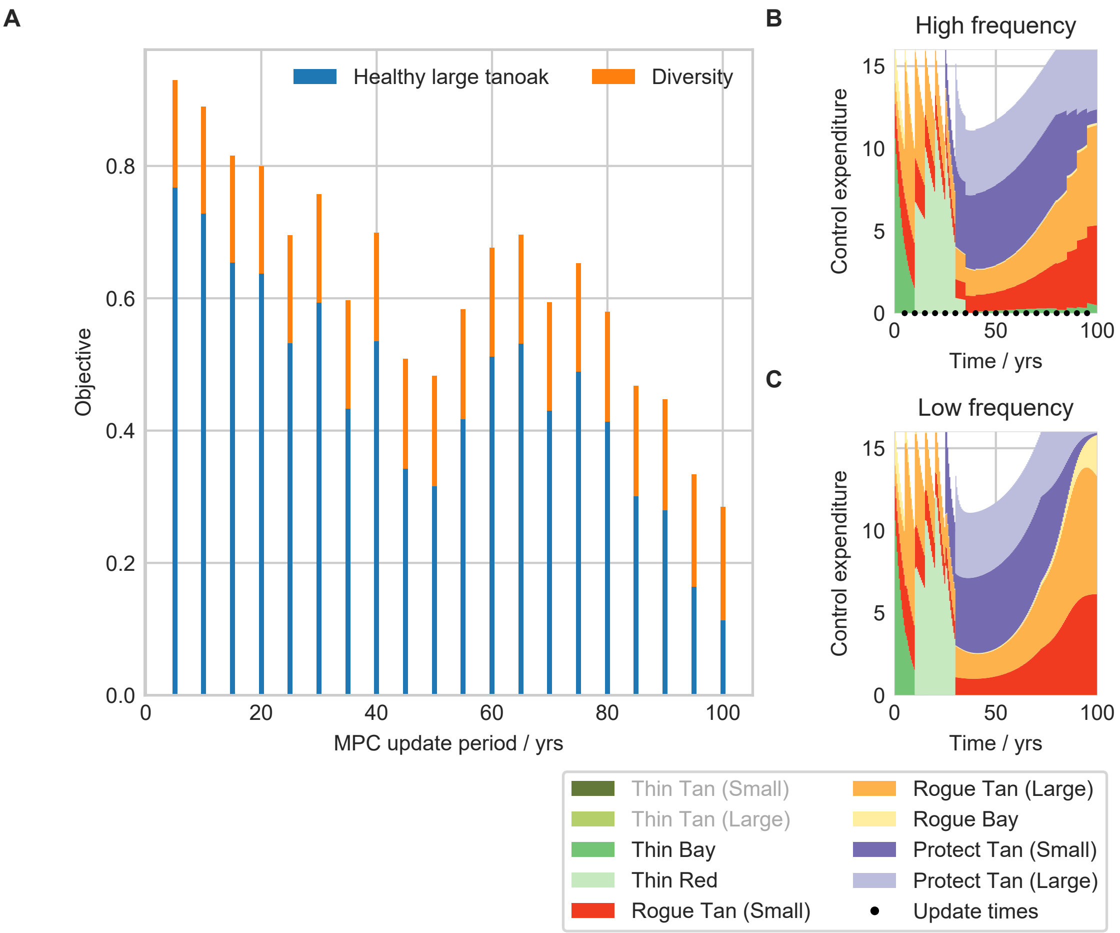


Figure 1: Effect of the MPC update period on control performance. A: the objective value as a function of how often the MPC re-optimises control. Reducing the time until the next update generally improves control, although update periods of around 50 years perform worse than expected. This is because these periods do not update close to the late outbreak, and therefore miss the unexpected increase in infection. B and C show the control allocations for update periods of 5 and 100 years respectively. The low frequency control here corresponds to open-loop control. The high frequency control results in more continued thinning than in the low frequency control. The update times are shown as black circles in B.

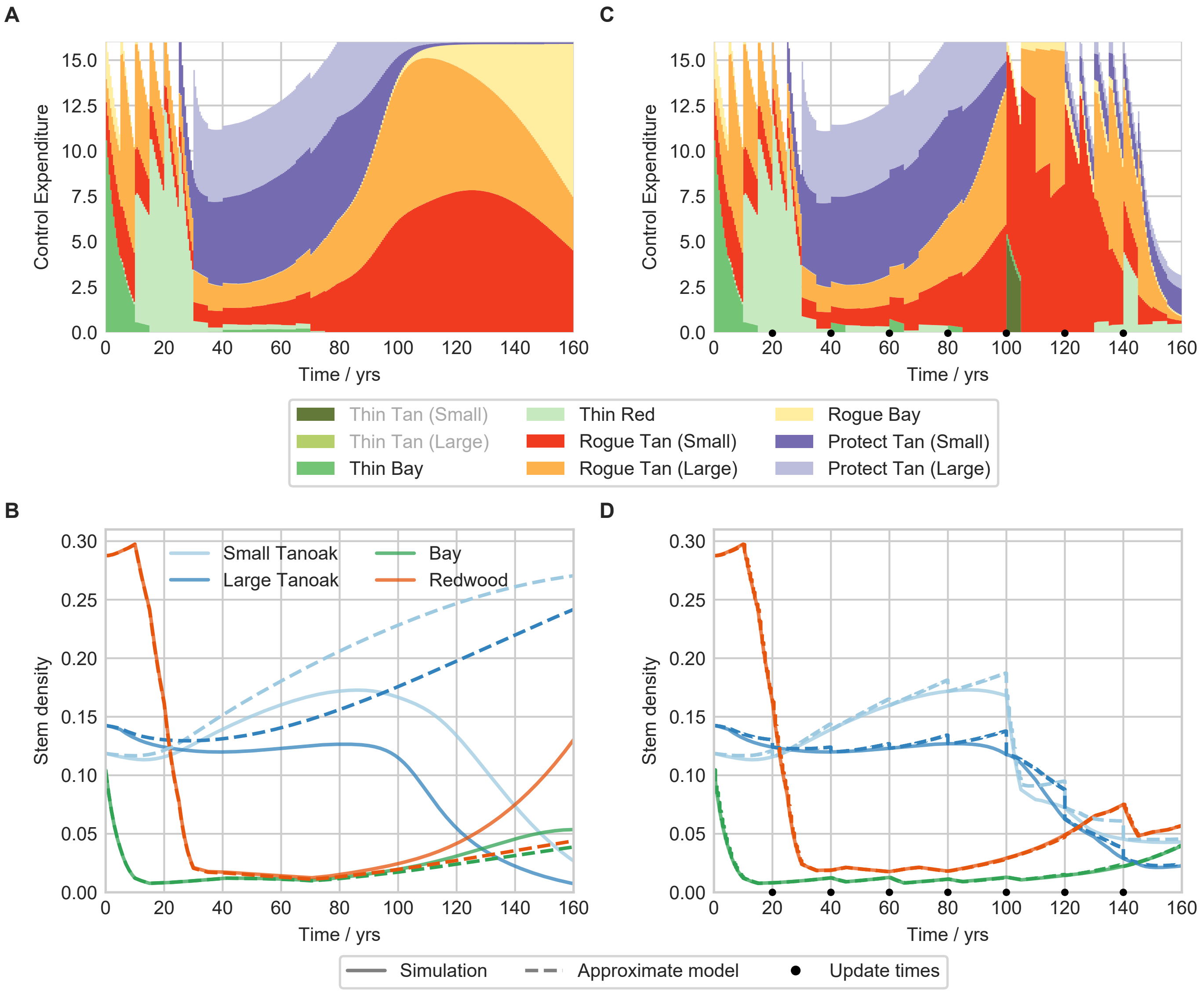


Figure 2: Open-loop and MPC control strategies, and host dynamics, using a time horizon of 160 years. A and B are the control strategy and host dynamics for open-loop, C and D show the same for MPC. More tanoak is retained using MPC, and the population has stabilised. MPC slows down the tanoak decline seen using the open-loop framework.

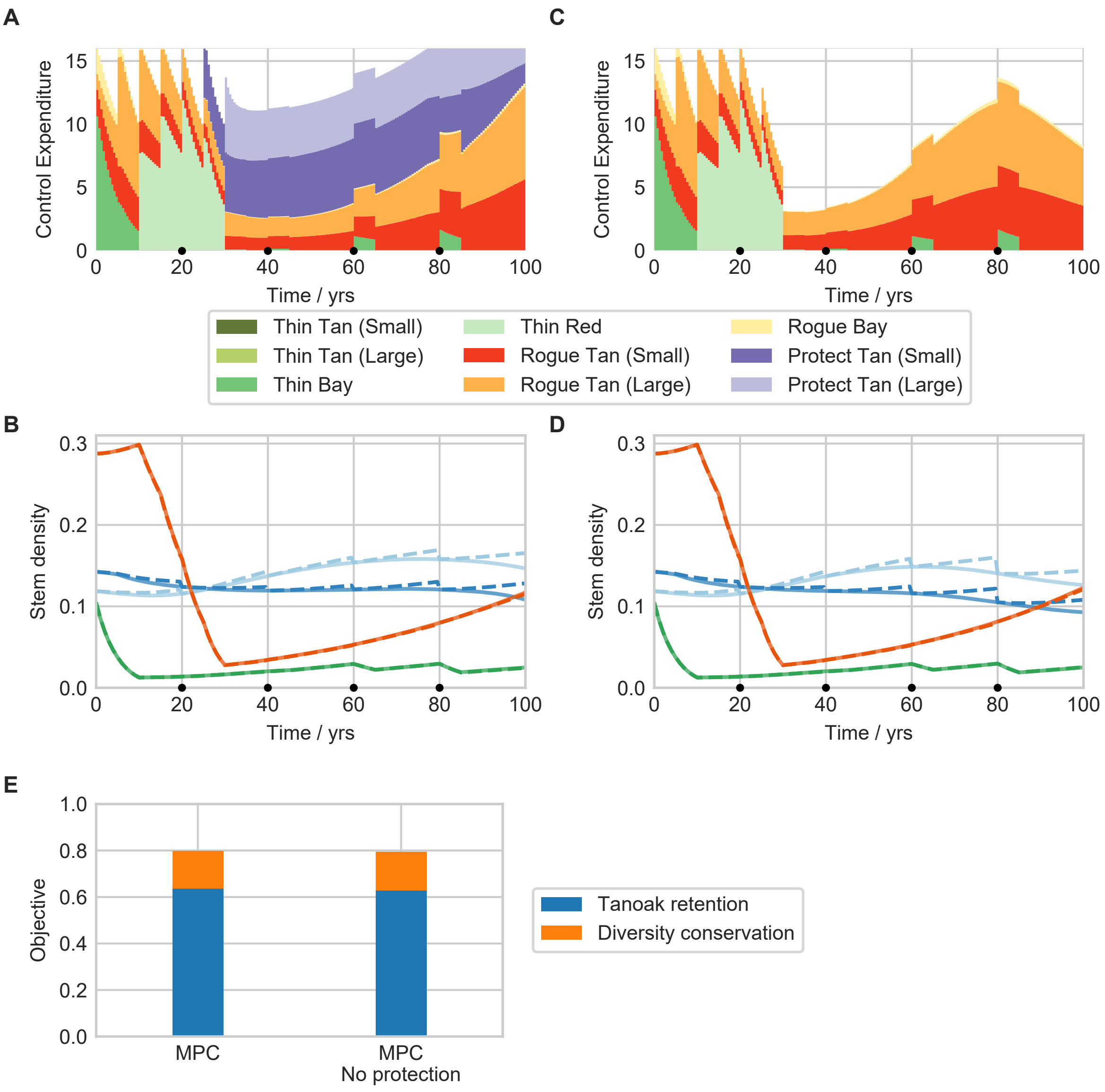


Figure 3: Under the MPC framework, applying protectant methods does not have a significant effect on the control performance. A and B show the control and host dynamics under the standard MPC framework. C and D show the same but where the protectant method is not applied. Note that control expenditure, in particular for roguing, is not the same since the number of hosts has changed. E: The objective function for MPC with and without protectant methods. The total objective values are 0.7996 (4 s.f.) and 0.7938 (4 s.f.) with and without protectant application respectively.

©The Authors under the terms of the Creative Commons Attribution License http://creativecommons.org/licenses/by/3.0/, which permits unrestricted use, provided the original author and source are credited.
