## Supplementary material for "Ongoing surveillance protects tanoak whilst conserving biodiversity: applying optimal control theory to a spatial simulation model of sudden oak death": S4 Text

### Supporting Information S4 Text

E.H. Bussell & N.J. Cunniffe

#### S4 Parameter uncertainty

In Figure 5C–5D in the main text, reproduced for convenience in Figure 1 below, 4 scenarios were highlighted to show the differences between open-loop and MPC under different parameter sets.

**Scenario 1** Open-loop performs badly, MPC is significantly better.

**Scenario 2** Average open-loop performance, moderate improvement by using MPC.

**Scenario 3** Good performance using open-loop, marginal decrease in performance using MPC

**Scenario 4** Good performance using open-loop, marginal increase in performance using MPC

Figures 2–5 and the captions describe the situation in each scenario, explaining what drives the differences in performance.

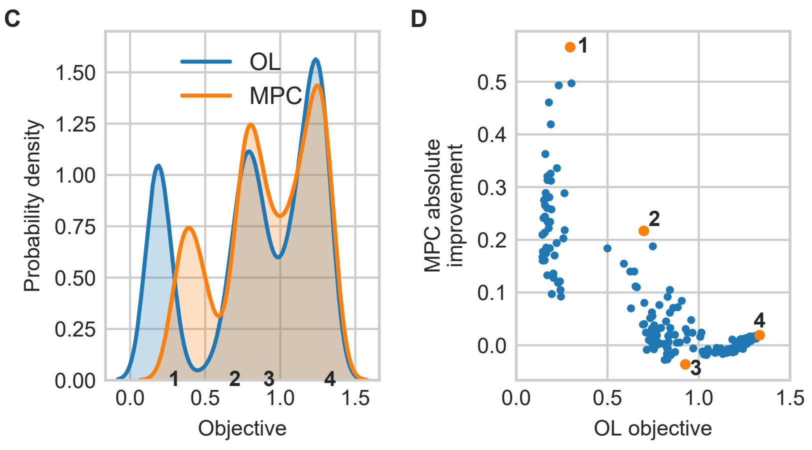


Figure 1: Repeat of Figure 5C–5D from the main text. C: The distribution of objective values using open-loop and MPC across 200 draws of simulation parameters. D: The absolute improvement of the MPC strategy over open-loop, as a function of the open-loop objective.

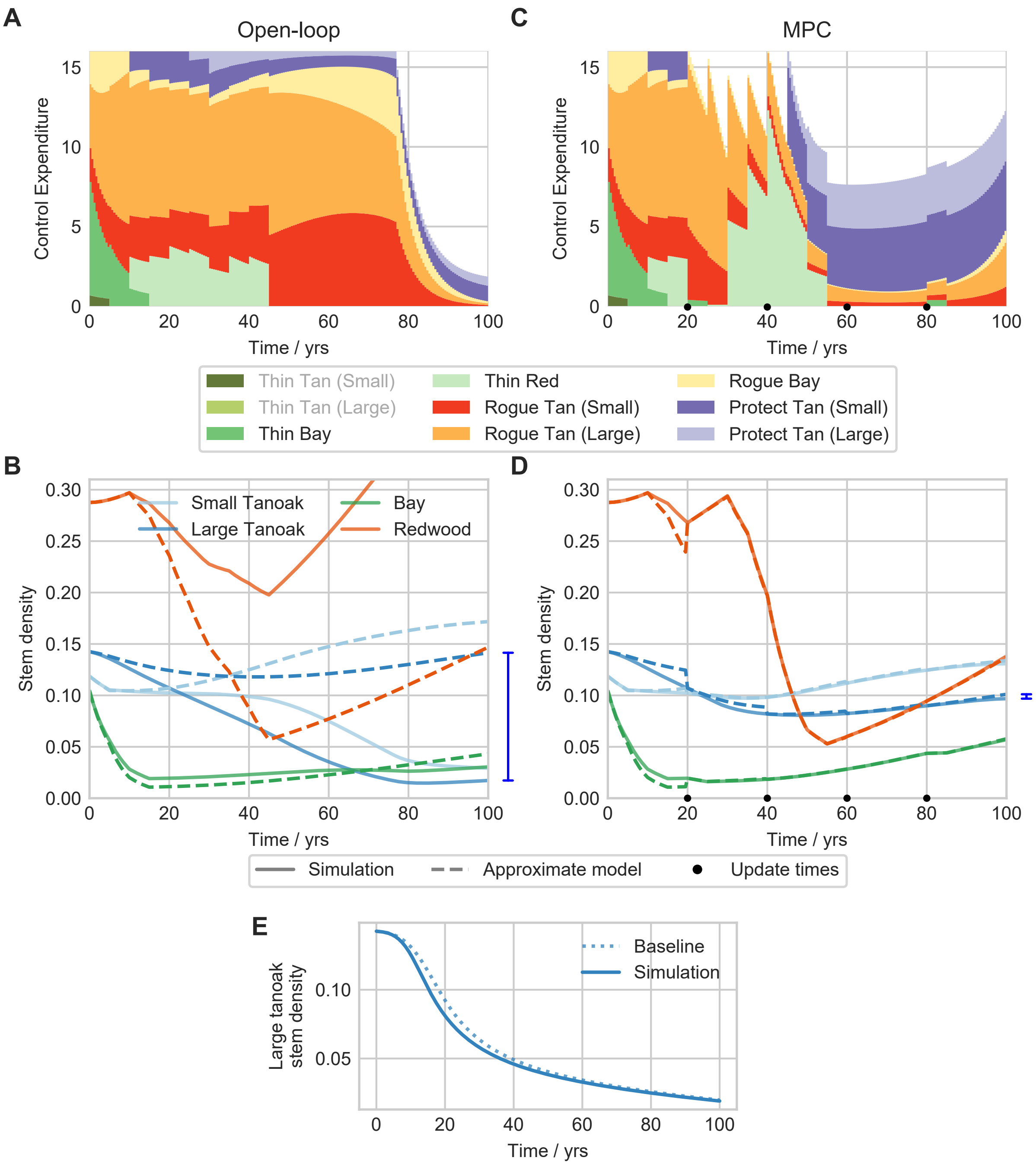


Figure 2: Scenario 1. For each scenario, the open-loop control strategy and host dynamics are shown in A and B, and the same for MPC in C and D. The blue bars to the right of B and D highlight the difference between the simulation and approximate models in the number of large tanoak at the final time. E shows the large tanoak dynamics when there is no control compared with the baseline parameter case. Here, open-loop performs poorly because the disease spreads quickly, leading to significant tanoak decline in the first 20–40 years. E shows decline is faster than in the baseline case. In MPC, the framework can respond to this early decline and keep the disease under control.

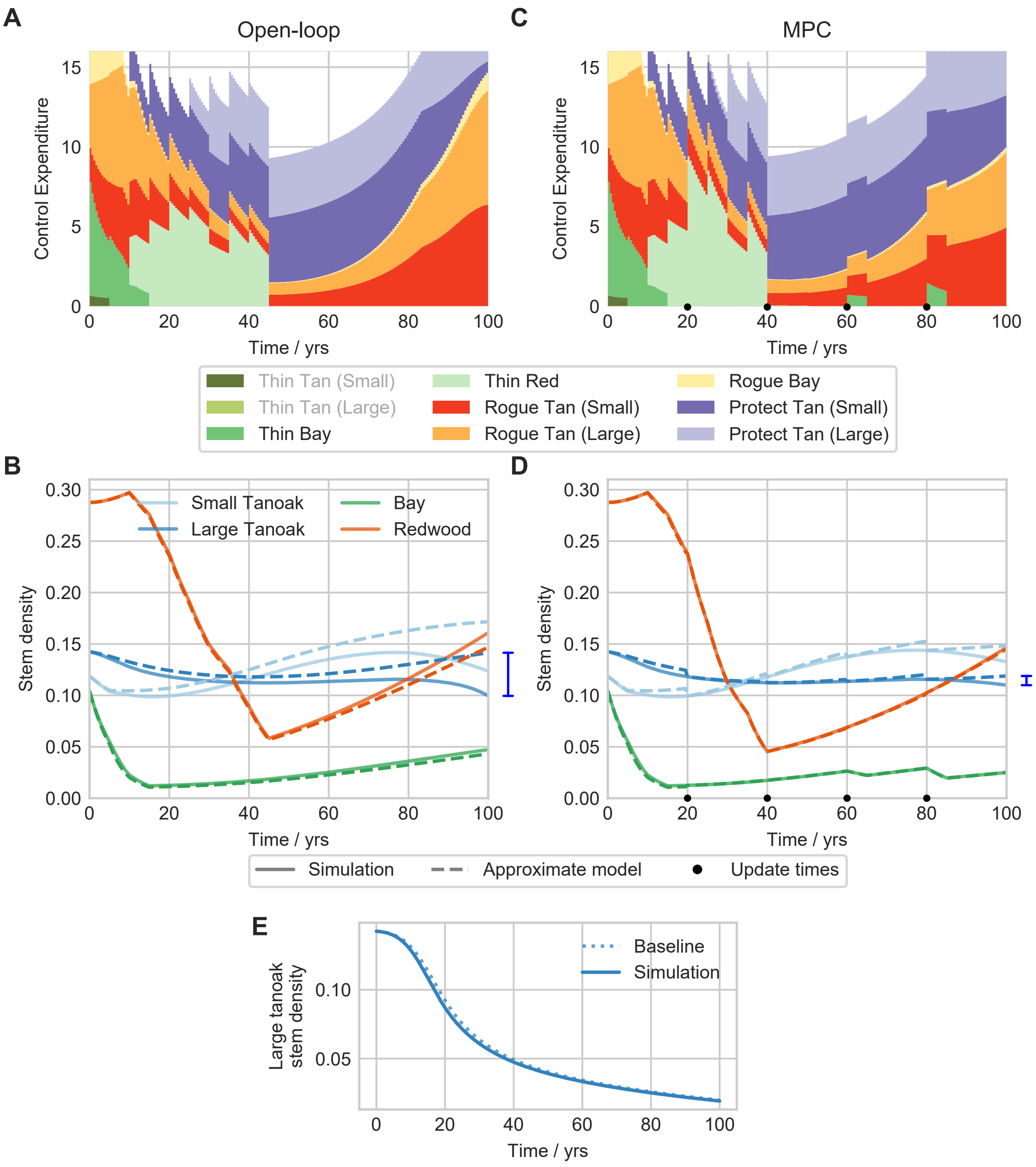


Figure 3: Scenario 2. Here, under open-loop the approximate model slowly degrades and leads to differences between the simulation and approximate models. The control is relatively effective, but is not informed by the correct simulation state. Under MPC the approximate model is kept much closer to the simulation, leading to more informed control and better performance. E: Tanoak decline under no control is similar to the baseline case.

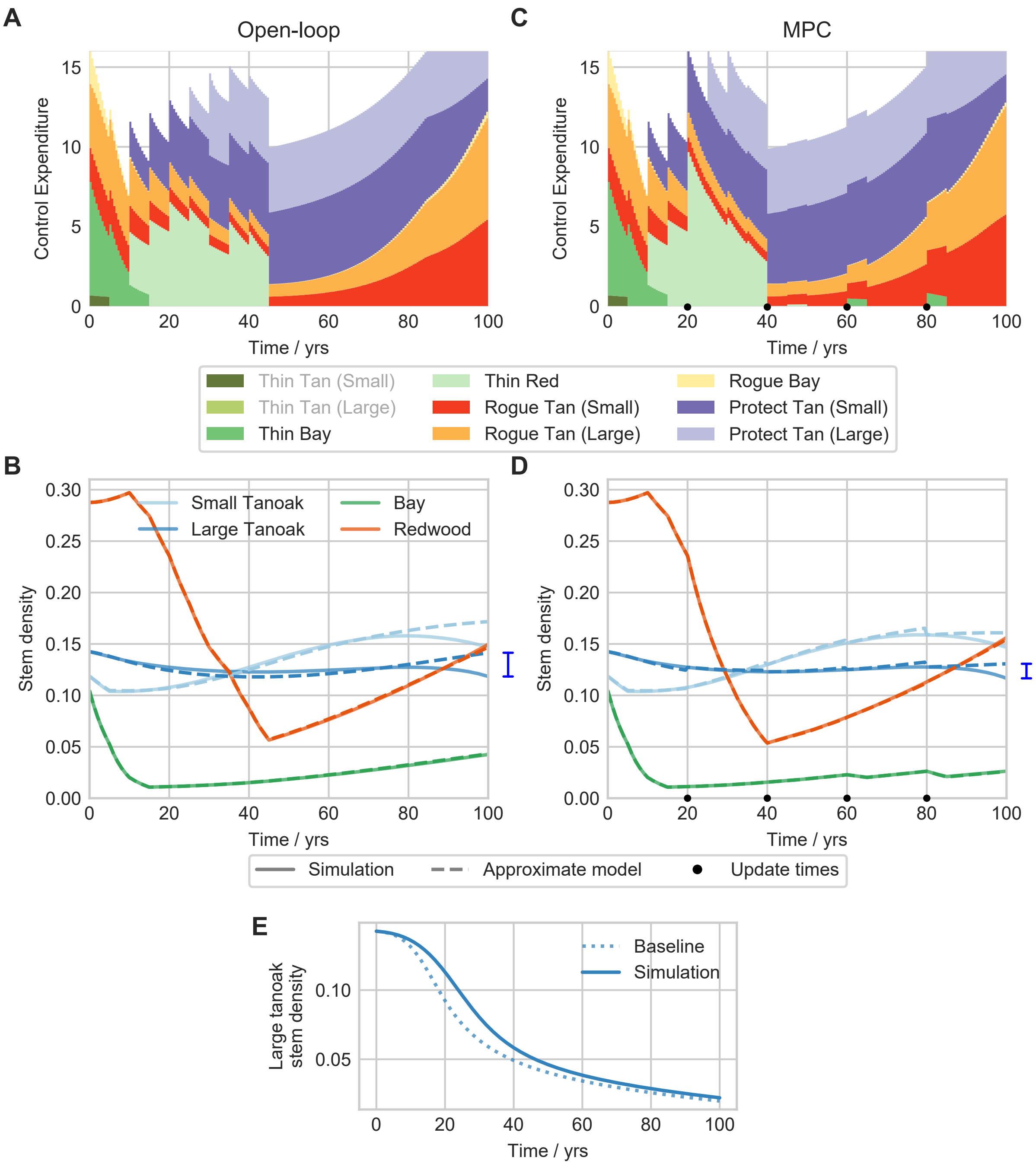


Figure 4: Scenario 3. The disease is slow to spread, and therefore relatively easy to control. The approximate model stays close to the simulation under both open-loop and MPC as there are only small amounts of disease spread. The different thinning regime under MPC leads to slightly worse retention of tanoak than under open-loop, but the difference is very small. E: Under no control, tanoak decline is slower than in the baseline case.

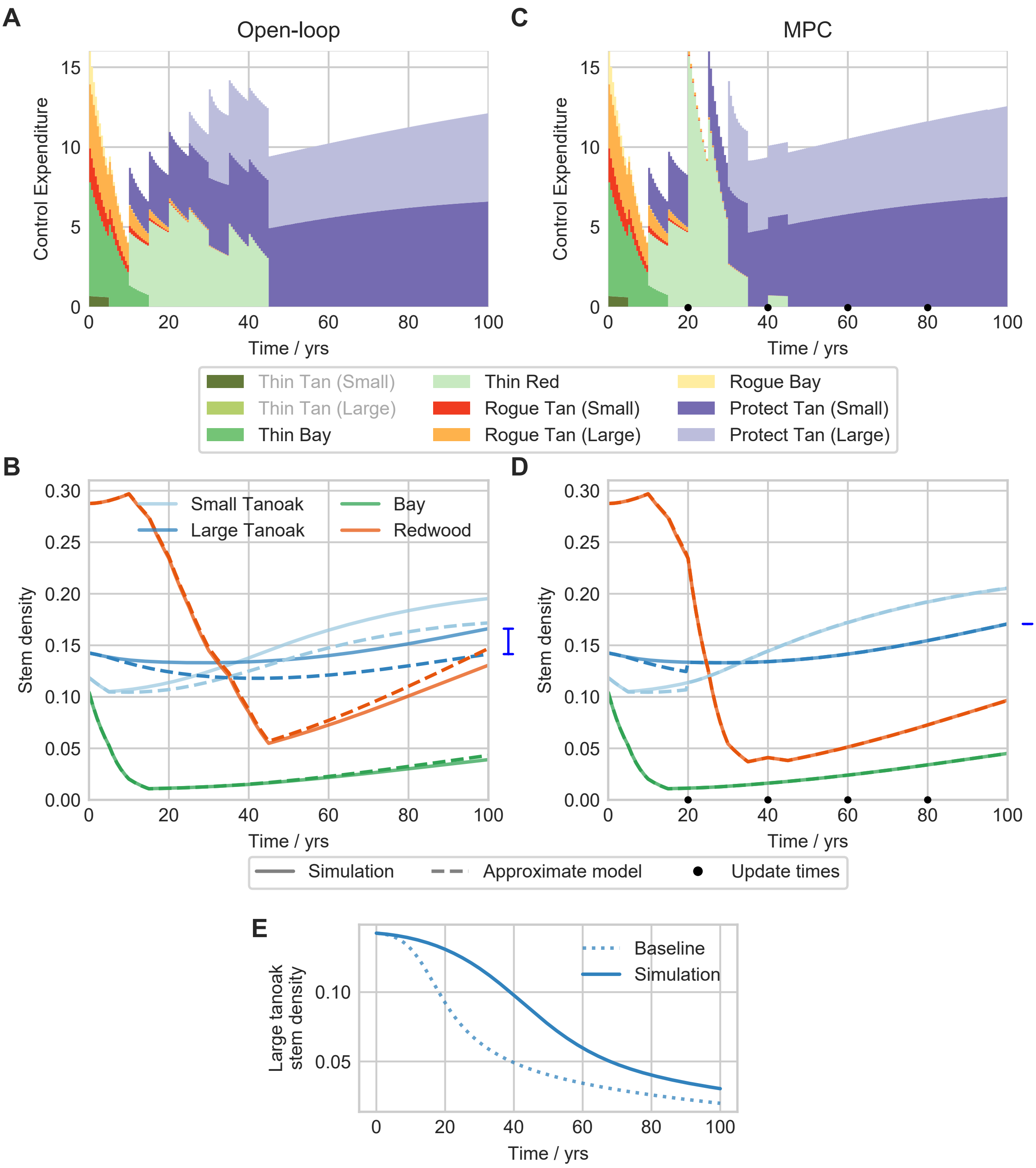


Figure 5: Scenario 4. The disease is very easy to control, leading to minimal roguing under both frameworks. The thinning of redwood under MPC is better informed after the update time at 20 years, and so promotes additional recovery of tanoak. Here both frameworks increase the size of the tanoak population above the pre-disease introduction level. E: Under no control, tanoak decline is much slower than in the baseline case.

©The Authors under the terms of the Creative Commons Attribution License http://creativecommons.org/licenses/by/3.0/, which permits unrestricted use, provided the original author and source are credited.
